# DeePhy: A Deep Learning Model to Reconstruct Phylogenetic Tree from Unaligned Nucleotide Sequences

**DOI:** 10.1101/2025.03.09.642239

**Authors:** Aritra Mahapatra, Jayanta Mukherjee

## Abstract

1. Inferring phylogenetic trees for a set of taxa is one of the primary objective in the evolutionary biology. Numerous approaches exist to reconstruct the phylogenetic trees by considering different biological data, such as DNA sequence, protein sequence, protein-protein interaction graph, etc. However, each method has its own strengths and weaknesses. Till date, no existing method guarantees to determine true phylogenetic trees all the times. Various studies identified distinct branch length configurations where the existing methods are inefficient to infer the correct tree topologies.
2. Here, we propose a novel deep convolutional neural network (CNN)-based model, *DeePhy*, to reconstruct the phylogenetic trees from the unaligned sequences. The sequences are represented on a two-dimensional coordinate plane by utilizing a biological semantics-based mapping. Additionally, to assess the robustness of a method, here we also propose a novel bootstrapping technique to generate replicas from the unaligned sequences. We train the model on the triplet sequences, where the output is a triplet tree topology.
3. We show that the well-trained DeePhy outperforms the state-of-the-art methods in inferring triplet tree topology. We experiment DeePhy on data simulated under numerous critical conditions and various branch length configurations. We conduct the McNemar test for comparing the performance of DeePhy and the state-of-the-art methods. The results exhibit that DeePhy is significantly more accurate and remarkably robust in determining the triplet tree topologies for most of the cases than that of the conventional methods. Again, various comparison metrics show that DeePhy also outperforms the conventional methods in inferring trees. Finally, to analyze the performance of DeePhy on real biological dataset, we apply it on Gadiformes dataset. Reassuringly, DeePhy reconstructs the phylogenetic tree from real biological data with known or widely accepted topologies.
4. Although various practical challenges still need to be taken care of, the outcomes of our study suggest that the deep learning approaches be a successful endeavour in inferring the accurate phylogenetic trees.

## 1 Introduction

Computational biologists observe mutations in various homologous segments of DNA sequences while studying the phylogeny of different operational taxonomic units (OTUs) by exploiting molecular data (Felsenstein, 2003; Jones and Pevzner, 2004). This approach is sensitive to the selection of segments from the sequence (e.g. genes, coding segments, etc.). Multiple sequence alignment (MSA) process tries to align the same segments of a set of complete mitochondrial genome sequences. However, organization of the homologous regions within the mitochondrial sequence may get violated during the evolutionary processes when the genes change their order within the genome sequences or get fragmented (Bernt et al., 2013). Apart from that, MSA is a computationally expensive process. For these reasons, attempts have been made to bypass this process and derive the phylogenetic trees directly from the unaligned gene sequences. We refer to this approach as alignment-free method and the tree derived from the individual gene sequences is called the gene tree (Maddison, 1997). While a gene tree tells the story about the evolution of the corresponding gene, on the other hand, a species tree represents the evolutionary history of the species (Szöllöosi et al., 2014; Degnan and Rosenberg, 2009). In one category of the alignment-free methods, DNA (or protein) sequences are represented numerically and are placed on the multi-dimensional coordinate plane. Finally, the pairwise distances are obtained by comparing these numerical or graphical representations (Zielezinski et al., 2017; Haubold, 2013; Randíc et al., 2013). It is hypothesized that every organism carries some unique signatures within their gene (or protein) sequences (Langille et al., 2010). Exploiting signatures from the genome sequences and utilizing them in phylogenetic tree reconstruction is the key motivation behind the mathematical representation of a genome. In the numerical representations, the encoded genome sequences are considered as the signal (called *genome signal* ). Different signal-processing techniques extract meaningful unique signatures from it (Yin and Yau, 2015, 2017). Existing numerical representations are broadly grouped into two categories – fixed value-based mapping and biological semantics-based mapping (Mendizabal-Ruiz et al., 2017). The fixed value-based mapping employs arbitrary numerical values to each nucleotide, whereas the biological semantics-based mapping associates numerical values with the biochemical or biophysical properties of the nucleotides. It is observed that the biological semantics-based mapping with multi-dimensional representations is more accurate than that of the fixed value-based mapping for the biological experiments (Mendizabal-Ruiz et al., 2017). The total number of possible topologies of a rooted phylogenetic tree for *n* number of OTUs is (2*n* − 3)!! which is equivalent to [1n (2k − 3) (Felsenstein, 1978b). Determining the topology from a set of sequences (both nucleotide and protein) is considered as an NP-hard problem (Roch, 2006). However, none of the methods are able to determine the true topology of a phylogenetic tree for a set of OTUs all the times.

Supervised machine learning approaches have been widely used in the biological studies for predicting diseases and classifying species based on their taxonomy groups (Das and Mitra, 2021; Mahapatra and Mukherjee, 2020; Tarca et al., 2007). Deep neural architectures provide powerful models for performing these tasks (Goodfellow et al., 2016). In supervised learning, for a given multivariate observation *x* there is a label *y* associated with it. By learning a model, it seeks to determine a function *f* (*x*) that predicts *y*. This is achieved through a process called *training*, whereas the *x* associated with *y* is called the training set. By utilizing the big data the deep learning networks show very attractive performances in different domains including biology (Das and Mitra, 2021; Najafabadi et al., 2015; Ching et al., 2018; Tarca et al., 2007). Unlike a conventional machine learning technique, the deep learning networks extract meaningful feature vectors from the data to achieve *f* (*x*). Thus they do not require any predefined feature vectors. Deep learning models require a huge amount of data for training the model. However, availability of true phylogenetic data is very limited. Therefore, providing the true relationships among a set of OTUs to the model becomes a challenging task in the phylogeny domain (Zou et al., 2019; Lemmon and Lemmon, 2013).

This creates the main bottleneck in applying sequence data-based deep learning models in the phylogenetic studies. A few proposed deep learning-based approaches derive phylogenetic trees from both nucleotide and protein sequence data. Surprisingly, all these techniques adopted the fixed value-based representations (Suvorov et al., 2020; Zou et al., 2019; Zheng et al., 2019; Kasperski and Kasperska, 2016). Apart from that, most of these methods provided an unrooted tree from a set of sequences (Suvorov et al., 2020; Zou et al., 2019), whereas the rooted topology of a phylogenetic tree contains more precise evolutionary information.

In this study, we adopt a biological semantics-based mapping to represent the mitochondrial genome sequences in a two dimensional coordinate space. We propose a novel convolutional neural network (CNN)-based model, called *DeePhy*, which is able to reconstruct the phylogenetic tree from a set of nucleotide sequences. We show that DeePhy can be trained to extract meaningful signatures from the 2*D* numerical representations which can be used to accurately reconstruct the rooted phylogenetic tree topologies. We map the computational problem into a classification task, where DeePhy is trained to classify the input three numerical representations into three possible topologies. We conduct in depth analyses to understand the performance of DeePhy on the datasets simulated under various possible circumstances. We find that DeePhy is highly accurate and its performance is fairly robust on the simulated data. We also exhibit its performance on the real biological dataset. Various computed performance metrics show that DeePhy outperforms the conventional phylogenetic tree reconstruction methods. In the subsequent sections, we describe our methodology and the performance of DeePhy in details. We also describe the strength and weakness of DeePhy, the real-life challenges, and the possible future extensions of the proposed deep learning-based approach in phylogenetic studies.

## 2 Materials and Methods

DeePhy is an alignment-free method that can reconstruct the phylogenetic tree from the nucleotide sequences of the OTUs. The nucleotide sequences are numerically represented on a two-dimensional coordinate plane. This numerical representation is called Genomic Footprint (GFP) of the corresponding nucleotide sequence (Mahapatra and Mukherjee, 2020, 2021b). DeePhy takes this GFP of the nucleotide sequence of each OTU as the input and trains a novel deep learning model to derive the phylogenetic tree. In the next section, we first describe the GFP of a nucleotide sequence briefly. Then, in the subsequent sections, we discuss in detail about the proposed deep learning model.

### 2.1 Genomic FootPrint (GFP)

The GFP of a nucleotide sequence is a locus on a two dimensional coordinate plane. There are three different structural groups of nucleotides based on i) the number of nitrogen rings (purine and pyrimidine), ii) the strength of inter-chained Hydrogen-bonded interactions (strong and weak H-bond), and iii) the kind of tautomerism (amino and keto).

Let *S* = {*s*_1_*, s*_2_*, . . . , s_n_*}, be a nucleotide sequence of *n* nucleotides, where *s_i_* be the the *i^th^* nucleotide of *S*. In information theory, self-information refers to the degree of certainty carried by the occurrence of an event (Chen, 2016). The self-information of an event *x* is derived from the probability of the occurrence of *x*. To compute the probability of a codon the second order Markov chain is considered where, we compute the probability of the occurrence of a nucleotide given two previous nucleotides. Hence, the probability of occurrence of a codon (*s_i−_*_2_*s_i−_*_1_*s_i_*) can be represented as,

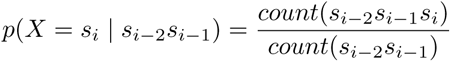

where the number of occurrence of an event *x* is denoted as *count*(*x*). Now, the probability of occurrence of a codon is used to determine its self-information. If the self-information of the *i^th^*codon, (*s_i−_*_2_*s_i−_*_1_*s_i_*), is represented as *I_i_*, then,

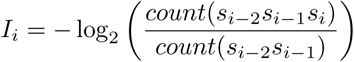

Similarly, the probability of the *i^th^* nucleotide of *S* is denoted as *r_s_*, and given by

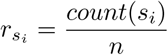

Since *s_i_* ∈ {*A, T, G, C*}, there are four probabilities correspond to four nucleotides, *A*, *T* , *G*, and *C*. They are denoted as *r_A_*, *r_T_* , *r_G_*, and *r_C_*, respectively.

If (*x_i_, y_i_*) denotes the coordinate of the *i^th^* nucleotide of the sequence *S*, then the values of *x_i_* and *y_i_* can be determined by utilizing the following set of rules, Case-1: Purine (R)/Pyrimidine (Y)

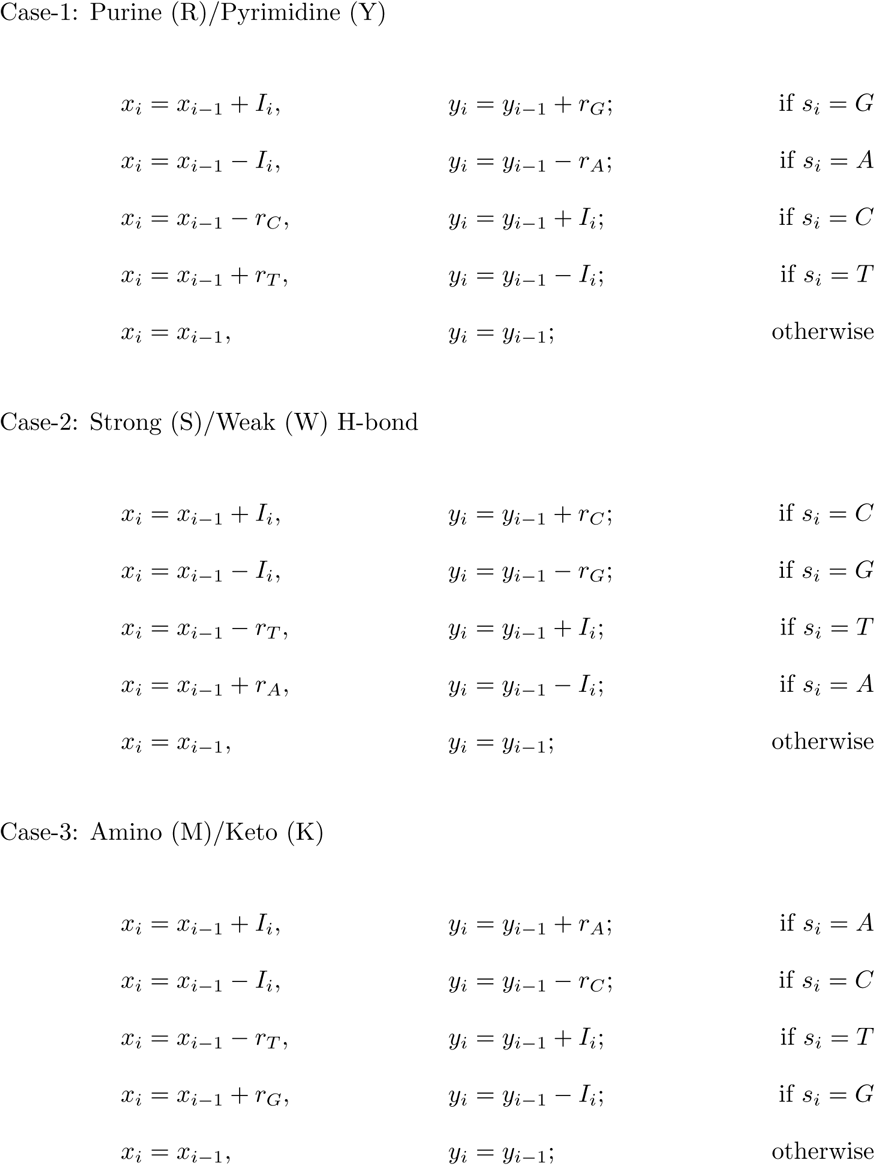

A recent study (Mahapatra and Mukherjee, 2021b) exhibited that among the three representa- tion schemes, the Amino-Keto representation captures more meaningful signatures from the genome sequences than that of the other representations. Hence, in this study, we consider the Amino-Keto representation scheme to represent a nucleotide sequence numerically on a 2*D* coordinate space. Our proposed model considers the GFPs of the sequences derived by utilizing the Amino-Keto representation scheme. In the following sections, we refer to this representation as “GFP” only.

### 2.2 Utilization of triplet

Deep learning approach is highly dependent on the data. However, in the phylogenetic study, the availability of true tree is very limited. To overcome this limitation, we utilize the triplet data in this study. A triplet tree, as shown in Fig. 1(a), is the smallest possible rooted binary tree which has three taxa or OTUs. A rooted phylogenetic tree of *n* number of leaf nodes can be decomposed into a set of number of triplet trees (Semple and Steel, 2003). An example is shown in Fig. 1(b) where the phylogenetic tree having five leaf nodes is decomposed into ten different triplet trees. On the other hand, we can also reconstruct a phylogenetic tree from the set of triplet trees. Our proposed method, DeePhy, applies a divide and conquer technique to reconstruct the phylogenetic tree. For *n* number of OTUs, DeePhy generates possible triplet trees which are utilized to reconstruct the phylogenetic tree by following a triplet tree amalgamation technique. We train the DeePhy model on a triplet dataset so that the model can successfully identify the triplet tree topology from the GFPs of three sequences. The simulation process of the triplet data is described in the next section.

**Figure 1:**
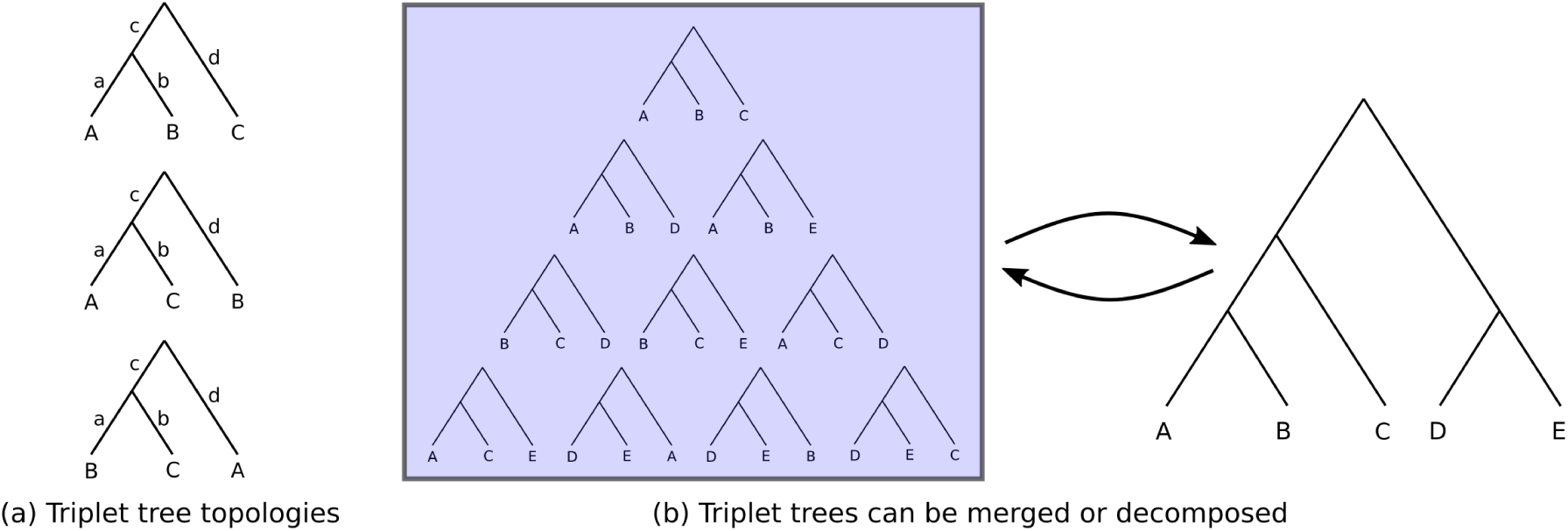
Illustration of triplet tree. (a) Three possible triplet tree topologies for three OTUs. (b) The decomposition of a rooted phylogenetic tree into a set of triplet trees. These set triplet trees can reconstruct the phylogenetic tree.

### 2.3 Triplet data simulation

The design of data plays a very crucial role in deep learning (Miotto et al., 2017; Najafabadi et al., 2015). For three OTUs, *A*, *B*, and *C*, there exist three different triplet tree topologies, such as ((*A, B*)*, C*), ((*A, C*)*, B*), and (*A,* (*B, C*)) in Newick notation format, which are also shown in Fig. 1(a). In this study, our data consists of two parts – a) triplet tree topology and b) nucleotide sequence of the OTUs of the respective triplet tree. The entire data simulation process follows five steps which are described below in details.

1. **Triplet simulation:** For this study, we simulate a large number of triplet trees consisting of the leaves, *A*, *B*, and *C*. We utilize a tool called Analysis of Phylogenetics and Evolution (APE) 5.0 (Paradis and Schliep, 2019; Paradis, 2012; Paradis et al., 2004) to simulate the triplet trees. During this simulation, APE also assigns the branch lengths of each triplet randomly from a uniform distribution. Since a triplet tree has four branches, it is expected of having a large varieties of branch-length configurations in simulated triplet trees.
2. **Eliminating criteria:** Among these different branch lengths we initially eliminate all the triplet trees having branch length greater than one. Now, for a triplet tree, there are two different relationships – sibling and non-sibling. For this study, we consider that the distance between any two sibling OTUs is less than the distance between two non-sibling OTUs. This case can be mathematically represented as, (*a*+*b*) *<* (*a*+*c*+*d*) and (*a*+*b*) *<* (*b*+*c*+*d*), where, *a*, *b*, *c*, and *d* are the four branch lengths and (*a* + *b*) is the distance between two siblings, as shown in Fig. 1(a). We discard those trees which do not follow both of these conditions.
3. **Uniform selection of triplet trees:** After eliminating the inconsistent triplet trees, we have a pool of triplet trees from where we prepare our dataset. In the deep learning method, an unbalanced training dataset also hampers performance of the models (Das and Mitra, 2021). To maintain the class balance in the dataset, we apply a novel technique to select the triplets. In this technique, we consider a four-dimensional grid structure, where each dimension refers to a specific branch of the triplet tree (please refer to Fig. 1(a)). Now, each dimension contains a certain number of partitions of the same size. Each partition refers to a specific range of lengths of the corresponding branch. In this study, we select the resolution of the grid as 0.1. Since the branch lengths of the triplet trees may vary from zero to one, there are ten partitions at each dimension. Therefore, within the grid structure, there are total 10^4^ cells, where each cell contains the triplet trees having the branch lengths within a specific ranges. After eliminating the triplets based on the criteria discussed earlier, there are only 5, 995 number of cells which can contain any triplet trees. Finally, we randomly choose equal number of triplet trees from each of the selected cells. This technique ensures consistency in the branch length configuration within our dataset.
4. **Sequence simulation:** After selecting the triplet trees, our next objective is to simulate the sequences of the corresponding OTUs of each triplet tree for which we utilize a tool called INDELible (Fletcher and Yang, 2009). By considering a tree topology along with its branch length, INDELible generates the nucleotide sequences of the corresponding OTUs by simulating insertions, deletions, and substitutions. During simulation, this tool also takes a nucleotide substitution model into account. The debate on the nucleotide substitution model in the phylogenetic study also indicates the requirement of further research in order to develop more realistic model for nucleotide substitution (Abadi et al., 2019; Hoff et al., 2016; Arenas, 2015; Shapiro et al., 2006; Posada and Crandall, 2001). In this study, we consider two widely accepted nucleotide substitution models – Jukes-Cantor (JC) model (Jukes and Cantor, 1969) and General Time-Reversible (GTR) model (Rodriguez et al., 1990; Tavaŕe, 1986) and simulate the sequences by considering these models separately. The JC model is the simplest nucleotide substitution model which assumes that the substitution of a nucleotide with any other nucleotide occurs with equal probability. Whereas, the GTR model is possibly the most general neutral, independent, finite-sites, time-reversible model. GTR model requires six substitution rates along with the frequencies of four nucleotides as the parameters. These four nucleotide frequencies are selected randomly from a four-category Dirichlet distribution with *α* = 30; whereas, the substitution rates are selected randomly from a uniform distribution, U(0, 3). For the insertion and deletion, we choose a fixed rate of 0.01 for both the operations. Finally, we set the length distributions in insertion and deletion from the Zipfian Distribution with *a* = 1.5, where the maximum length of insertion and deletion is also set as 5% of the total length of the simulated sequence. Since DeePhy considers unaligned sequences, the lengths of the sequences may vary from each other.
5. **Numerical representation:** Now, we have a set of three simulated sequences of the corresponding OTUs of a triplet tree. Finally, we derive the GFP of each sequence by utilizing the Amino-Keto representation scheme and consider these GFPs for training, validation, and testing purposes of our proposed deep learning model. The GFP of a sequence contains same number of 2*D* points as the number of nucleotides present in it. Hence, the lengths of the GFPs of different sequences may be different in the designed dataset.

For this study, we randomly choose 6.5 × 10^5^ number of triplet trees by taking almost equal number of samples from each of the selected cells of the grid. Then, we simulate the sequence of the corresponding OTUs of each triplet tree by considering each nucleotide substitution model separately. During simulating the sequences, we choose the length as 1, 000 bp. However, since we consider the set of unaligned sequences, the lengths of the sequences may vary from each other.

Hence, for each nucleotide substitution model, we simulate 6.5×10^5^ set of sequences, where each set contains the sequences of an approximate length of 1, 000 bp of three OTUs of the respective triplet tree. Then we derive the GFPs of the sequences which are considered as the input for DeePhy.

### 2.4 Model architecture and Training

Here, we propose a deep learning model which can identify the topology of a triplet tree for a set of three OTUs based on their corresponding GFPs. For three OTUs, *A*, *B*, and *C*, there exist three different triplet tree topologies - ((*A, B*)*, C*), (*A,* (*B, C*)), and ((*A, C*)*, B*). We can also represent the same three topologies based on the outgroup OTUs. So, we consider this problem as a three-class classification problem, where each triplet tree topology is considered as a separate class. For a triplet tree, the sibling OTUs are considered as the ingroup OTUs, whereas the third one is the outgroup. Hence, alternatively, we can also say that each class corresponds to an outgroup OTU among the three OTUs. We prepare the input by representing the three GFPs as a matrix where each row corresponds to a OTU. Since GFP of the nucleotide sequence is a 2*D* representation, the values corresponding to *x*-axis and *y*-axis are placed at the separate channels of the input matrix. So, for *L* length of triplet sequences, a 2 × 3 × *L* dimension of input is provided to our model.

Now, we propose a novel convolutional neural network (CNN)-based model (Goodfellow et al., 2016) to extract the features from the input matrix. Based on the feature vector the model also selects the class for the set of three inputs. The lengths of a nucleotide sequence and its corresponding GFP are same. Hence, the GFPs derived from different sequences vary in sizes. Since convolutional layer takes input of same length, we consider the longest GFP of the dataset and extend the smaller GFPs by padding with (0, 0) coordinate at the end. A deep learning model is tuned by various hyper-parameters and there are no fixed rules to select their proper values. Hence, we apply a grid search technique to choose the hyper-parameters. For this technique, we randomly select 100 sets of test datasets independently and each test dataset contains 3 × 10^4^ triplet data (combination of a triplet tree and the sequences of the corresponding OTUs). We select the model undergoing least validation loss. We apply the model on 100 test datasets separately and compute the mean and the standard deviation of the accuracies. We finally use these statistics to determine the hyper-parameters of the model in the search space. Details of the grid search process is shown in the **Supplementary S1**.

Our proposed model, DeePhy, consists of five convolutional layers with the ReLU activation (Nair and Hinton, 2010) for each layer. Interestingly, for this study, it is observed that average pooling layer provides better performance than that of the maximum pooling layer which is more commonly used in extracting the features from input data. Accordingly, we apply the average-pooling operation with the size of 1 × 3 over the output of each convolutional layer. The total number of filters for the first and the last convolutional layers are set as 64. Whereas, for three intermediate convolutional layers, this number is set as 128. We set the filter size of the first convolutional layer as 2 × 3. For the other convolutional layers, we set this size as 1 × 3. Additionally, for all the convolutional layers, we also set the stride as 1 × 1. The CNN layers are utilized to extract the features from the input matrix. After the stack of convolutional layers there exist one fully connected layer having 64 nodes with ReLU activation followed by another fully connected layer having three output nodes corresponding to the three possible tree topology classes. The fully connected layers take the responsibility of classifying data based on the feature vector extracted by the CNNs. To avoid the generalization error and to normalize the output of the previous layers, we use batch normalization technique (Ioffe and Szegedy, 2015) after each convolutional layer and also after the first fully connected layer. Apart from that, to reduce the model overfitting, we also use dropout regularization (Baldi and Sadowski, 2013, 2014) with a rate of 0.10 at each convolutional layer only. Our proposed network is shown in Fig. 2.

**Figure 2:**
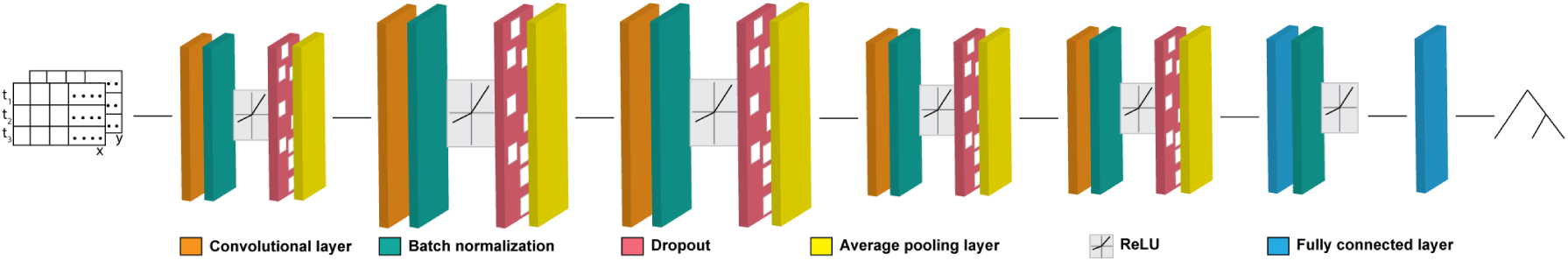
The architecture of DeePhy. The network input is the GFPs of three sequences, rep- resented as t_1_, t_2_, and t_3_. The dimensions (*x* and *y*) of the GFPs are provided as the separate channels at the first CNN. The network output is a class corresponding to a triplet topology.

We have simulated 6.5 × 10^5^ sequences by considering JC and GTR nucleotide substitution models separately and derived their respective GFPs. Each set of data consists of three GFPs and the respective class representing a triplet tree topology. The labels of the classes are one-hot encoded where the sibling and outgroup OTUs are represented by 0 and 1, respectively. Considering each nucleotide substitution model separately, we select 6 × 10^5^ data randomly and use them to train the model. From the rest of the data, again we choose 3×10^4^ data randomly and use it as the validation dataset. To observe the effect of different nucleotide substitution models in the phylogenetic study, we train our model with the GFPs derived from each nucleotide substitution model separately. We use the Adam optimizer (Kingma and Ba, 2015) with an initial learning rate of 0.00001. The cross-entropy loss is used to measure the error of each prediction. We decay the learning rate by halving the rate after each iteration if no improvement is found in the current validation loss over the best prior epoch. To avoid model overfitting, the training procedure runs for a number of epochs determined by an early stopping criteria and a patience value. The early stopping criteria is a threshold which indicates a significant improvement in each epoch. Whereas, the number of consecutive epochs the improvement is below the early stopping criteria is the patience value. We stop the training process if there are less than 0.00001 drops of the current validation loss over the previous one for consecutive ten epochs. From Fig. 3, it is observed that the performance of the training procedure is stable as it consistently achieves similar accuracy and loss on both the training and validation sets at each epoch. We implement DeePhy in Python 3.6 using PyTorch version 1.5.1.

**Figure 3:**
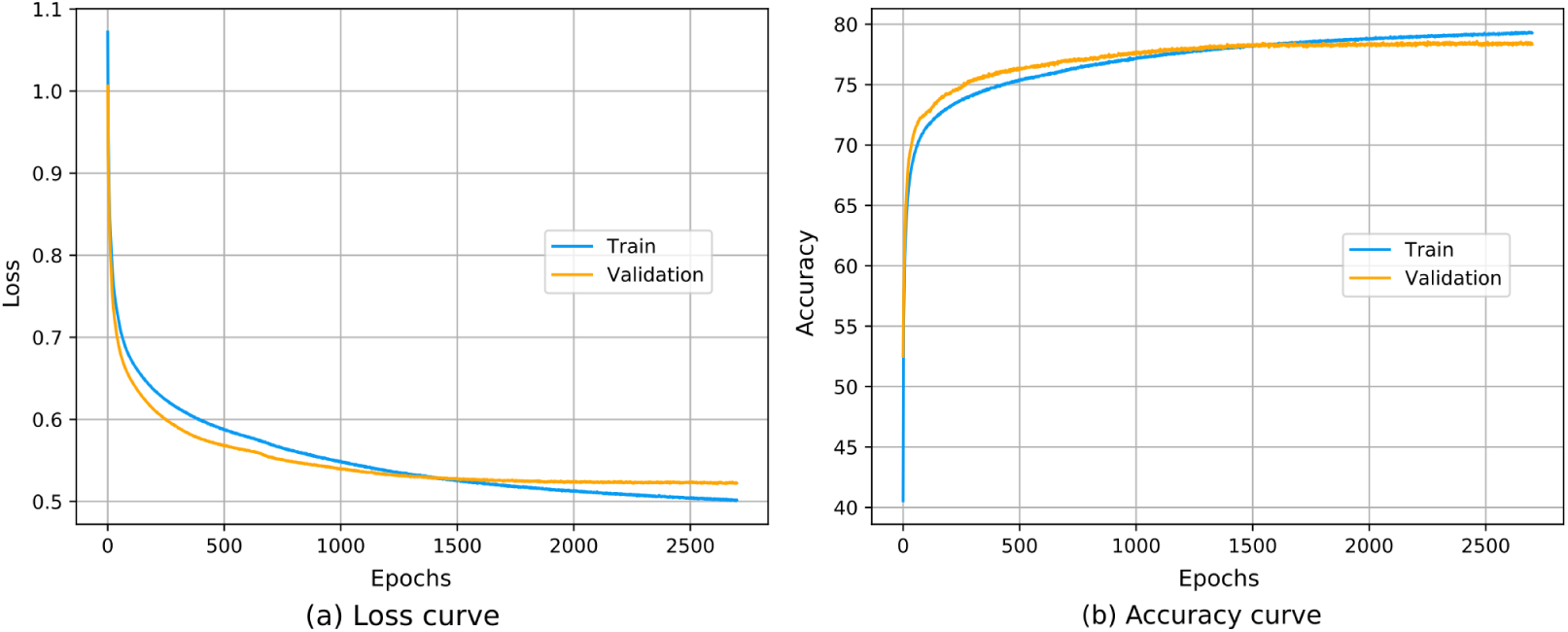
The (a) loss values and (b) accuracies of the training and validation sets at each epoch.

### 2.5 Testing procedures

For testing purpose we keep the test dataset separated from the training procedure. For a detailed performance analysis, we prepare 100 independent test datasets, where each set contains × 10^4^ triplet data. We compare the performance of DeePhy with the conventional methods, namely Neighbour Joining (NJ), Maximum Parsimony (MP), Maximum Likelihood (ML), and Bayesian Inference (BI) using our test datasets. Since all these conventional methods generate unrooted trees, they require at least four OTUs for deriving the trees. Therefore, these methods are not applicable on the triplet data. In order to evaluate their performance, we add a dummy sequence of length 1, 000 bp with each triplet sequence. During simulation of the dummy sequence, we select the nucleotide at each position with equal probability. So there is a very high chance for the dummy sequence to be the outgroup among four OTUs. Now, the conventional methods provide an unrooted tree, whereas, DeePhy provides a rooted tree as output. In order to compare the performance of DeePhy with the conventional methods, we need to convert both the output trees into rooted trees. Since the dummy OTU is the outgroup among the four OTUs, we apply the outgroup method (Kinene et al., 2016; Wheeler, 1990; Tarrıo et al., 2000; Hess and De Moraes Russo, 2007; Boykin et al., 2010) to root the unrooted tree. This method generates the rooted trees with four OTUs. Finally, we prune the dummy OTU from the derived trees. We utilize the tool MUSCLE (version 3.8.31) (Edgar, 2004a,b) for the multiple sequence alignment (MSA) and MEGAX software (Kumar et al., 2018) for deriving the trees from the MSAs by considering NJ, MP, and ML methods. Apart from that, we utilize 3.2.6 version of MrBayes software (Ronquist et al., 2012; Ronquist and Huelsenbeck, 2003) to derive the trees from the MSAs by using the Bayesian inference method. The commands we use to execute these methods are provided in **Supplementary S2**.

### 2.6 DeePhy on longer sequences

Conventionally, the size (or dimension) of the test data of a trained CNN model should match the size of the training data. In the phylogenetic study, to the best of our knowledge, all existing deep learning models considered smaller length of data to train their models. However, in the real-world, the whole mitochondrial sequences (mtDNAs) are quite longer than that of the training dataset. Due to this reason, the existing deep learning models are not applicable on the mtDNAs. Similarly, in this study, we simulate the training dataset which consists of sequences having nearly 1, 000 number of base pairs. During the training process we consider the longest sequence among the training dataset, which is 1, 356 bp. Since the GFP of a sequence contains the same number of 2*D* coordinate points as the length of the sequence, the longest GFP in our training dataset has 1, 356 number of 2*D* points. We extend the smaller GFPs in the training dataset by padding with (0, 0) coordinates at the end. Hence, the DeePhy model is applicable directly on the input data of 2 × 3 × 1356 dimension. There is a standard and well established approach that trains model on the dataset which contains mitochondrial sequences of same lengths (Suvorov et al., 2020). However, this extension requires a significantly large amount of resource and time. Here, we apply DeePhy on the longer sequences by using a majority-consensus approach to understand that whether we can develop a trained scalable deep learning model. In this approach, we make overlapping equal length partitions of the GFP of each mtDNA where each partition contains 1, 356 number of 2*D* coordinate points. For triplet data, we consider each set of partitions and apply the trained DeePhy model on it. This gives a class label based on the partitions. Similarly, for each set of partition, DeePhy provides the corresponding class labels. Finally, majority consensus is considered as the final class label of the triplet of whole mtDNAs. This method is illustrated in Fig. 4.

**Figure 4:**
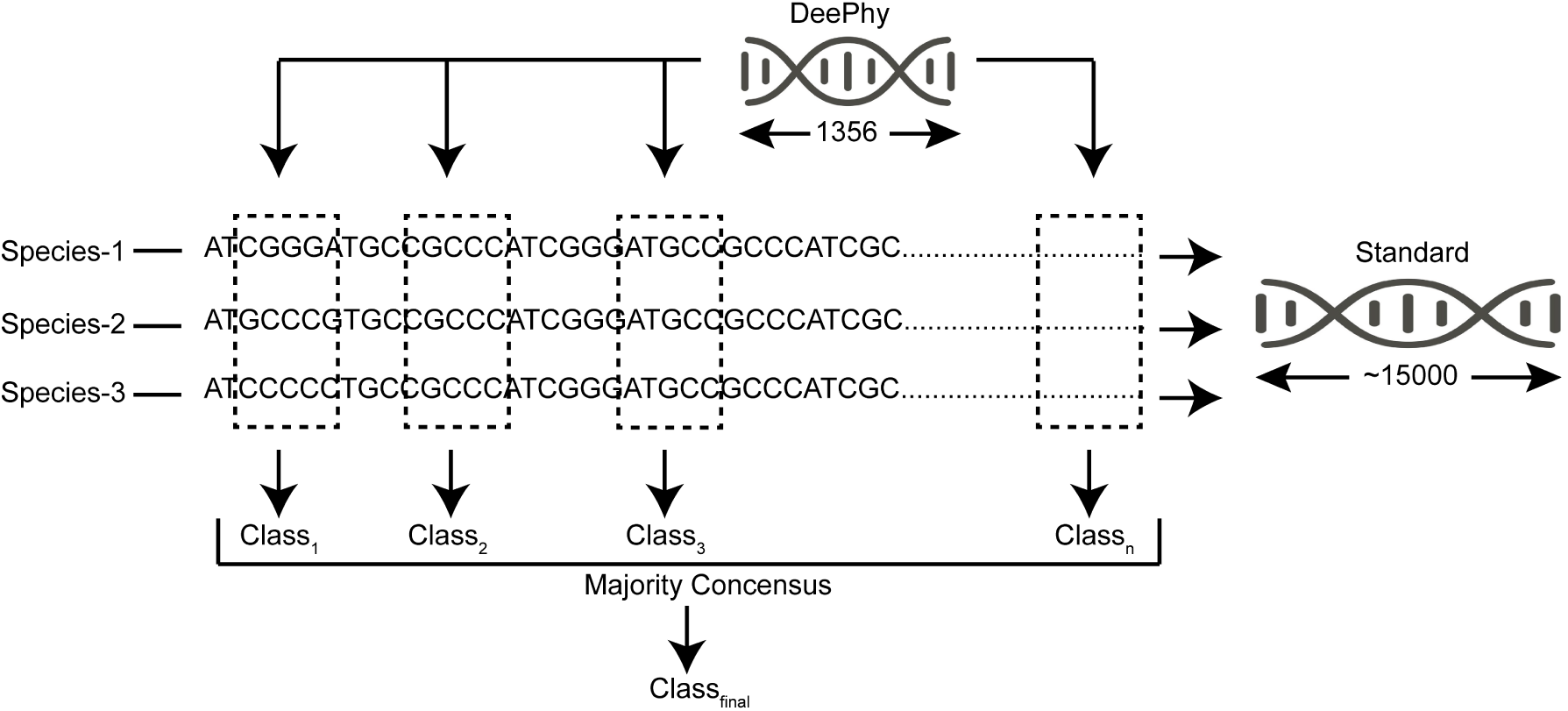
An illustration of the technique to determine the class for the long sequences using the trained DeePhy model. The longer sequences are split into overlapping partitions of length 1, 356 which is the length of the data in our training dataset.

### 2.7 Final tree construction

DeePhy follows divide and conquer approach to reconstruct a phylogenetic tree. For a *n* number of OTUs, DeePhy derives the possible triplet tree topologies. In the end, the final tree is constructed from the triplet trees by utilizing a triplet-based supertree reconstruction method, called SuperTriplets (Ranwez et al., 2010).

## 3 Results and Discussions

### 3.1 Bootstrapping

Generally, the existing phylogeny studies consider OTUs from different populations where an OTU is considered as a representative of the corresponding population. However, two represen- tatives from the same population do not have the same genome sequences. All the methods for inferring the phylogenetic trees from the sequences depend on the data. Therefore, if a method is sensitive to the sequence, the trees derived from the method may vary when different representatives are selected from the same population. This is not desired in phylogenetic study. In this context, the bootstrap technique determines the robustness of a method by applying a resampling technique to estimate statistics on a population (Lahiri, 2006). A bootstrap value of a clade indicates the percentage of occurrence of the same clade within the trees derived from the bootstrap replicas.

The conventional method of bootstrapping (Felsenstein, 1985; Efron et al., 1996) considers the aligned sequences to resample and replicate. As we are developing an alignment-free method for reconstruction of the phylogenetic tree, the conventional bootstrapping method may not be applicable for this case. The main motivation of bootstrapping is to generate the population from a single nucleotide sequence statistically. Various studies on both microbial and macro-organisms observed that the average intraspecific genetic variation of the mitochondrial genome is within 1% (Liu et al., 2020; Jarczak et al., 2019; Yang et al., 2014; Wang et al., 2008; Levy et al., 2007; Santamaria et al., 2007). So here we propose a bootstrapping technique which considers the genetic variance of a sample space within 1%. For that we apply both insertion, deletion and mutation at each location with an equal probability of 0.33%, so that the total variation of the sequence becomes 1%. For mutation operation at a location, we replace the nucleotide with a new one which we select in an unbiased manner. We generate 100 replicas using the proposed bootstrapping method and construct trees from each set of bootstrapped sequences using the methods as discussed in the previous sections. Felsenstein’s bootstrapping method (Felsenstein, 1985) assesses robustness of the phylogenetic trees using the presence and absence of clades. For large scale genomics, Felsenstein’s bootstrap is not efficient as it is inclined to produce low bootstrap support (Lemoine et al., 2018). So here we apply a modified version of Felsenstein’s bootstrapping technique, where the presence of a clade is quantified using the transfer distance (Lemoine et al., 2018). The transfer distance (Charon et al., 2006) is the minimum number of changes required to transform one partition to other. We compute the bootstrap score of each clade by utilizing the tool BOOSTER (Lemoine et al., 2018). In the following sections, we derive the bootstrap replicas by using our proposed method. We compute the bootstrap scores of the trees constructed by various methods by utilizing the online version of BOOSTER tool (url: https://booster.pasteur.fr/new).

### 3.2 Overall performance

For assessing the performance of DeePhy over all the branch length configurations, we simulate the test dataset consisting of 3 × 10^4^ triplet trees. We consider an equal number of random trees from each of 5, 995 different branch-length subspaces. We simulate the unaligned sequence of each OTU having length of nearly 1, 000 bp. To observe the consistency in the performance of DeePhy, we simulate 100 independent sets of such test dataset. To compare the performance of DeePhy with the existing methods, we apply NJ, MP, ML, and BI methods on the same test datasets by utilizing a dummy sequence and root the derived unrooted trees by applying the outgroup method, as we have discussed earlier. It is observed that, for 12.29% (with a standard deviation of 2.34) of the data , MP method fails to derive the trees from the respective quartet sequences. For computing the accuracy of MP method, we discard these undetermined cases. For 100 sets of test datasets, the average accuracies of NJ, MP, ML, BI, and DeePhy are 69.32% (s.d. 0.785), 72.44% (s.d. 0.759), 71.21% (s.d. 0.787), 72.22% (s.d. 0.550), and 78.21% (s.d. 0.437), respectively. To analyze the performance of DeePhy over other standard tree estimation methods closely, we conduct the McNemar test (Yang et al., 2010; Fagerland et al., 2013) whose *p*-values are less than 1×10*^−^*^20^. This phenomenon strongly indicates that DeePhy is significantly more accurate than the other methods, like NJ, MP, ML, and BI. For 100 sets of test datasets, Fig. 5(a) also shows that the accuracy of DeePhy is consistently higher than the existing methods for all the test datasets.

**Figure 5:**
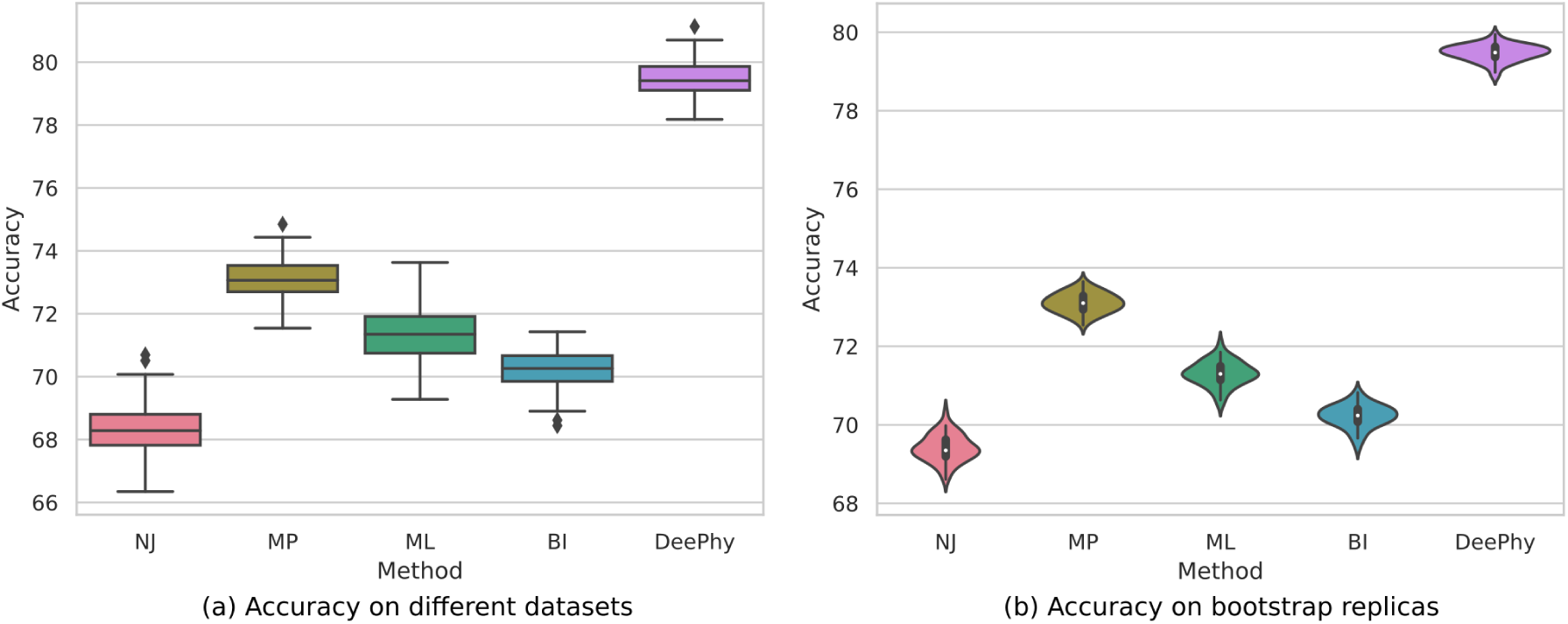
Overall performance of DeePhy and other existing methods on (a) 100 independent test datasets, (b) 100 bootstrap replicas of a test dataset. derive the tree by considering NJ, MP, ML, and BI methods, as discussed earlier. We also apply DeePhy on the bootstrap replicas. For each set of replica we compute the average accuracy. For 100 sets of bootstrap replicas, the average accuracies of NJ, MP, ML, BI, and DeePhy are 69.40% (s.d. 0.344), 73.11% (s.d. 0.25), 71.37% (s.d. 0.312), 70.22% (s.d. 0.292), and 78.82% (s.d. 0.281), respectively. Hence, from the bootstrapping, we can state that the robustness of DeePhy is comparable with the other existing methods. Please refer to Fig. 5(b). However, DeePhy shows more accuracy also in the bootstrap replicas of the entire branch-length space. In the next section, we conduct further studies by narrowing the branch-length space to observe the performance of the methods on various branch-length subspaces separately.

To assess the robustness of the methods, we select one of the test datasets randomly and derive 100 sets of bootstrap replicas of it. Since, DeePhy considers unaligned sequences, we apply our proposed bootstrapping method which generates the replicas from the unaligned sequences. To execute the existing methods such as NJ, MP, ML, and BI, on the bootstrap replicas we add a dummy sequence with each of the triplet sequences. After that, we align the quartet data and

### 3.3 Performance on different branch-length zones

There exist various regions in the branch-length parameter space where the performance of the existing methods is generally low (Suvorov et al., 2020; Bergsten, 2005; Roch et al., 2018; Felsenstein, 1978a). This bias is called as long-branch attraction (LBA) (Bergsten, 2005; Roch et al., 2018). For the unrooted tree, depending on the configurations, the regions in the branch-length parameter space effected by the LBA bias are termed as Felsenstein zone (Huelsenbeck and Hillis, 1993), Farris zone (Siddall, 1998), Twisted Farris zone (McTavish et al., 2015), etc. Since this study focuses on the rooted tree, we consider different branch-length configurations to assess the performance of both DeePhy and the existing methods for the rooted tree topology. We partition the entire branch-length space into five categories – extra-short, short, medium, long, and extra-long branch-length zones. For the extra-short, short, and medium branch-length zones, the lengths of all the branches of a triplet tree are within the ranges of [0, 0.1), [0.1, 0.4), and [0.4, 0.6), respectively. Whereas, for the long and extra-long branch-length zones, the length of atleast one branch of a triplet tree lies between the ranges of [0.6, 0.9) and [0.9, 1), respectively. Our selected branch-length configurations also include some branch lengths which we may not encounter in the phylogenetic study with real datasets – for example, having very short branch length between two OTUs indicates the presence of very less or no meaningful evolutionary changes between them. However, for the detailed analysis of the strength and weakness of DeePhy, we test each method on five different branch-length subspaces, separately.

To observe the performance for each of the branch-length zone, we simulate 1, 000 triplets from each branch-length zone separately and simulate the unaligned sequences of the respective OTUs. During the simulation, we again set the lengths of the sequences as 1, 000. However, this does not guarantee to simulate the unaligned sequences of same length. We apply both DeePhy and the existing methods on the dataset. Since the existing methods are incapable for dealing with the triplet relationship, we utilize a dummy sequence to derive the tree from the triplet data as discussed earlier. To observe the robustness of the methods in each branch-length space, we bootstrap each dataset and generate 100 replicas from each.

We observe that MP methods fail to derive the trees from the quartet sequences for some of the data of the test dataset. While measuring the performance of the method, we discard those cases where the method fails to derive the trees. For the rest of the cases, we compute the accuracies of the method for each branch-length zone. Fig. 6(a) shows that in most of the five branch-length zones, DeePhy exhibits significantly higher accuracies than that of the conventional methods. For the extra large branch-length zone, the accuracy of DeePhy (73.20%) is comparable with the best performing method in this branch-length zone, MP (73.50%). For the extra small branch-length zone, the MP shows an accuracy of 71.40% which is a little higher than that of DeePhy (68.40%). However, it is observed that, MP method fails to derive the trees for 91.10% of the quartet data taken from the extra small branch-length zone (please refer to Fig. 6(g)). Similarly, MP again fails for 51.50% of the data which are selected from the small branch-length zone. For MP, the number of failure cases decreases for increasing branch lengths. For inferring the quartet tree, both the differences between the opposing long and short terminal branches and the length of the internal branch are considered as the most important factor causing inconsistency in the performance of maximum parsimony (Schulmeister, 2004). In the extra short and short branch-length zones, both of the differences are small, that may be the reason for which MP encounters a large number of failure in deriving trees in both of these branch-length zones. We observe that based on the bootstrap technique, for each branch-length zone, the robustness of the methods are comparable to each other. Please refer to Figs. 6(b-f).

**Figure 6:**
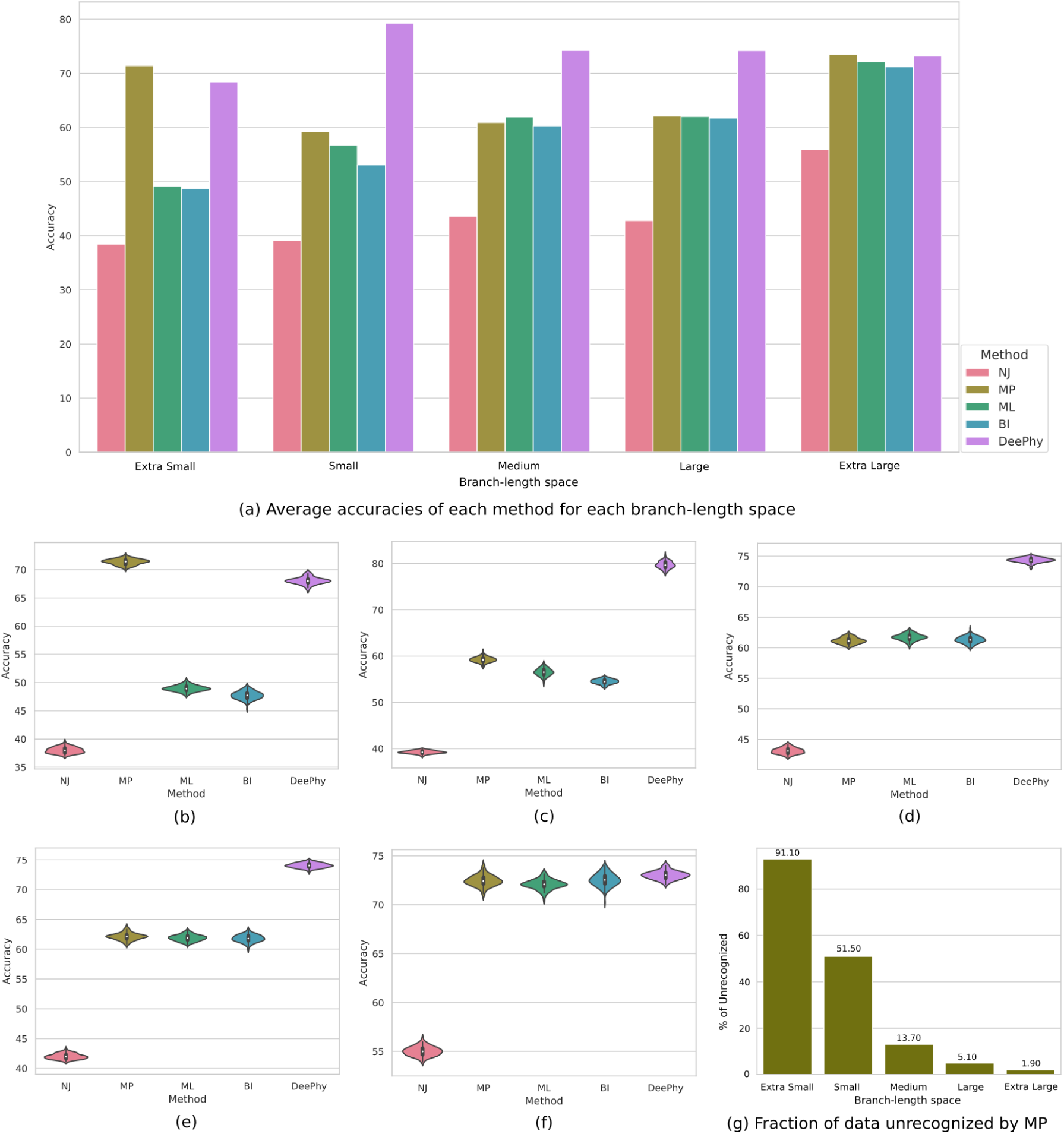
The performance of DeePhy and other existing methods for different branch-length zones. The triplets are selected from each branch-length space. For the existing methods, the same triplets are converted into quartet. (a) The average accuracies of each method for different branch-length spaces. The performances of the methods for 100 bootstrap replicas of the data selected from (b) extra small, (c) small, (d) medium, (e) large, and (f) extra large branch-length spaces. (g) The MP fails to derive quartet trees for some of the test data. The average number of cases for different branch-length spaces for which MP fails to derive the quartet trees. NJ = Neighbour Joining; MP = Maximum Parsimony; ML = Maximum Likelihood; BI = Bayesian Inference.

From this experiment, we observe that DeePhy outperforms all the conventional methods for every branch-length zones selected in this study. However, there are infinite number of triplet trees present in the entire branch-length space. For each of the branch-length zone, we have selected a subspace which is significantly smaller in size as related to the corresponding branch-length zone. For example, in this study, we have selected triplet trees having the branch lengths in the range of [0, 1]. If we discretize this range into 1000, then for the four branches of a triplet tree there will be 1000^4^ instances in the entire space. It is nearly impossible to experiment on each of the subspaces. So, we cannot eliminate the possibility that DeePhy is less efficient in some regions of the branch-length space. Hence, we are not over-ambitious about the performance of DeePhy. However, this performance may be improved by adding more data taken from the specific zones during the training process. This study requires extensive experiments on more subspaces of the branch-length space.

### 3.4 Performance on Longer sequences

We train DeePhy with the dataset where the length of the longest sequence is 1, 356 bp. Hence, the performance of DeePhy reaches the best level when the length of the input data is ≤ 1356 which we have shown in the previous experiments. However, in the real data, the mitochondrial sequences are quite longer than our training data. To make DeePhy applicable for the longer sequences, here, we propose a majority consensus-based technique where multiple class labels are derived from different segments of the triplet data. For this study, we again simulate 3 × 10^4^ random triplets and generate the sequence of each OTU by utilizing INDELible tool. During the execution of INDELible, we only change the length of the derived sequences as 15, 000 bp and keep the values of the other parameters same as earlier. Finally, we apply our trained DeePhy on the GFPs derived from the simulated sequences.

Since we have no idea about how much overlapping of the partition works best, we enumerate the process for various sizes of gaps in the overlapping. For an in depth analysis, we extend the gap larger than 1, 356 bp which leads to the non-overlapping partition. Since we adopt a majority consensus-based approach, there is a chance of having ties for some cases. Here, we denote the ties as the unclassified cases. We compute the accuracy based on the total number of data. From Fig. 7, we observe that the percentage of correctly identified triplet tree topologies does not change significantly by increasing size of gap. However, interestingly, the percentage of unclassified cases increases significantly with the increasing size of the gap. So, at this stage, we can say that since DeePhy is trained on short length data, it does not perform appropriately for the longer sequences by the majority-consensus approach. However, we believe this is a common problem for any deep learning-based method when the model is trained on shorter dimension of data and is applied on data having higher dimension. From this failure we can say that for the simulated sequences the signature is not equally distributed within a longer sequence as that of the shorter one. However, we believe this limitation can be overcome by training the DeePhy on the longer sequences.

**Figure 7:**
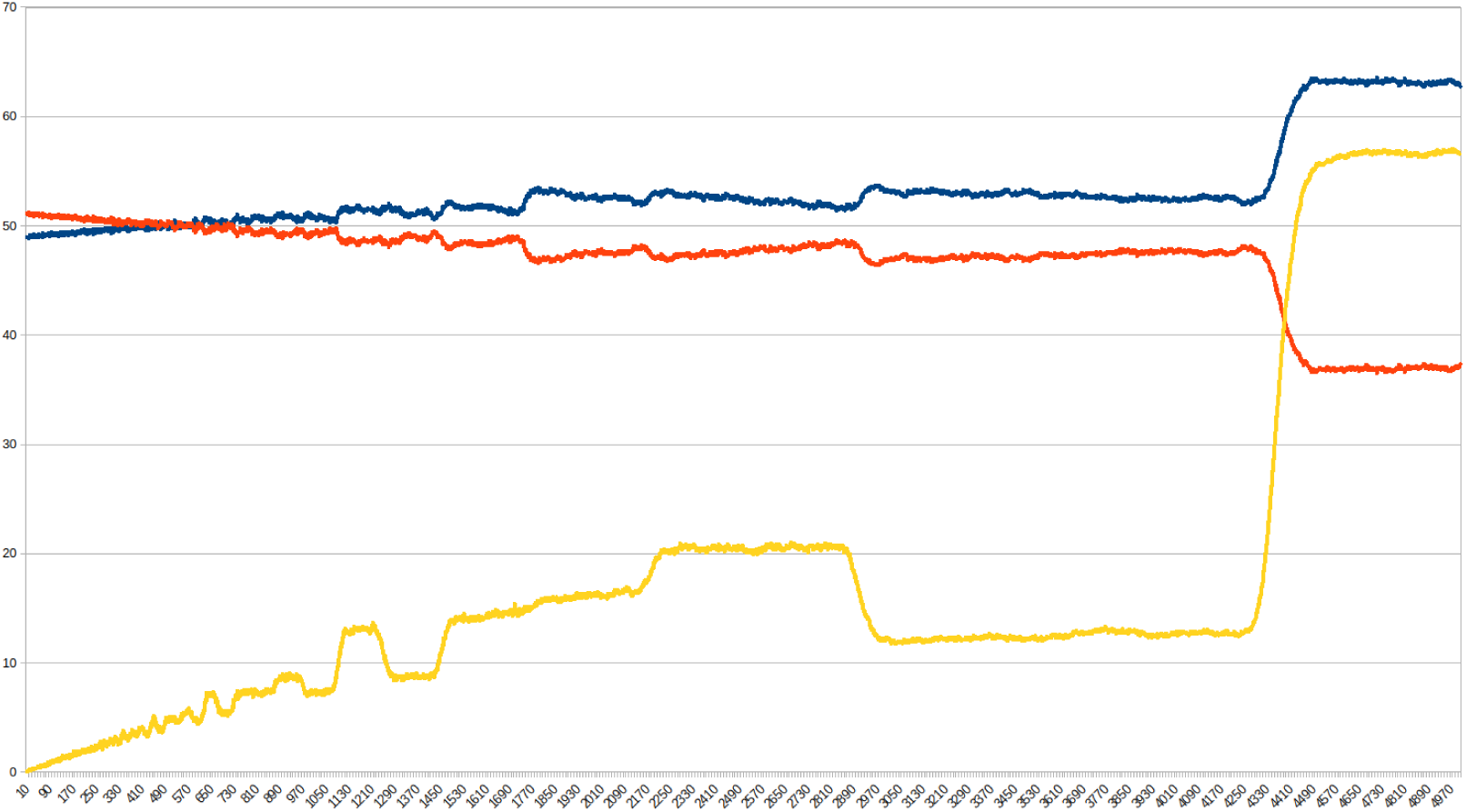
The percentages of correct (in blue), incorrect (in red), and unclassified (in yellow) cases for different size of gaps present in between two partitions.

### 3.5 Performance in deriving trees

The earlier sections have demonstrated the performance of DeePhy on various situations when the input is a triplet data. However, these studies do not exhibit the performance of DeePhy in deriving a complete phylogenetic tree. In this section, we apply DeePhy on a set of OTUs and finally form the complete phylogenetic tree. For this study, we consider phylogenetic trees having 10, 20, 30, 40, and 50 leaf nodes. For each case, we simulate 100 phylogenetic trees by considering coalescent theory. This ensures that for all the triplet subtrees of the simulated tree, the distance between two siblings is less than that of the non-siblings. During simulating the sequences, we set the length of the sequences as 1, 000 bp, therefore the lengths of the sequences simulated by utilizing INDELible automatically falls under 1, 356 bp. Finally, we apply both DeePhy and the other existing methods to derive the trees from each set of sequences. For *n* number of leaf nodes, DeePhy first makes all combinations of the OTUs and derives the triplet tree topology for each of the combinations. Finally, we utilize SuperTriplet tool to amalgam all triplet trees to produce the complete tree. To compute the accuracy of the derived tree, we consider the respective simulated tree as the reference tree and compute the Robinson-Fould (RF) (Robinson and Foulds, 1981), Matching Split (MS) (Bogdanowicz and Giaro, 2012), and Deformity Index (DI) (Mahapatra and Mukherjee, 2021a) scores between the derived trees and the reference trees. The conventional methods for comparing tree topologies, such as RF and MS, completely depend on the reference tree topology. So they may fail when both the reference and target trees contain different sets of OTUs. An alternative approach, Deformity Index, considers the existing hypotheses to compute the correctness of the target tree. In brief, DI takes a list of hypotheses regarding the phylogenetic relationships of the selected OTUs and compute the DI score of a target tree. In computing the DI, it takes each hypothesis separately and quantify the level of it’s support in the target tree. Finally, they are accumulated to derive the Deformity Index of the tree. This score represents the level of corroboration of the target tree with the list of hypotheses. The minimum value of DI is zero and it denotes that the target tree topology corroborates with all the available hypotheses completely. The larger DI score denotes less accurate tree with respect to the available hypotheses. This approach is considered as a more realistic measure especially for the biological datasets where the true reference tree topologies are unavailable. However, in this section, we compute the correctness of the derived trees by using both RF, MS, and DI scores.

It is observed that with respect to the reference trees, the RF, MS, and DI scores of the DeePhy derived trees are lower than that of the conventional methods for all the configurations of the leaf nodes. This phenomenon indicates that DeePhy simulates more accurate trees than that of the other standard methods like NJ, MP, ML, and BI. From Fig. 8, we also observe that the performance of DeePhy is better than that of the traditional methods for an increasing number of leaf nodes in the trees. So, till now, we have experimented both DeePhy and the conventional methods on the simulated data and after observing various comparison scores we can claim that DeePhy significantly outperforms the traditional methods for almost all the cases.

**Figure 8:**
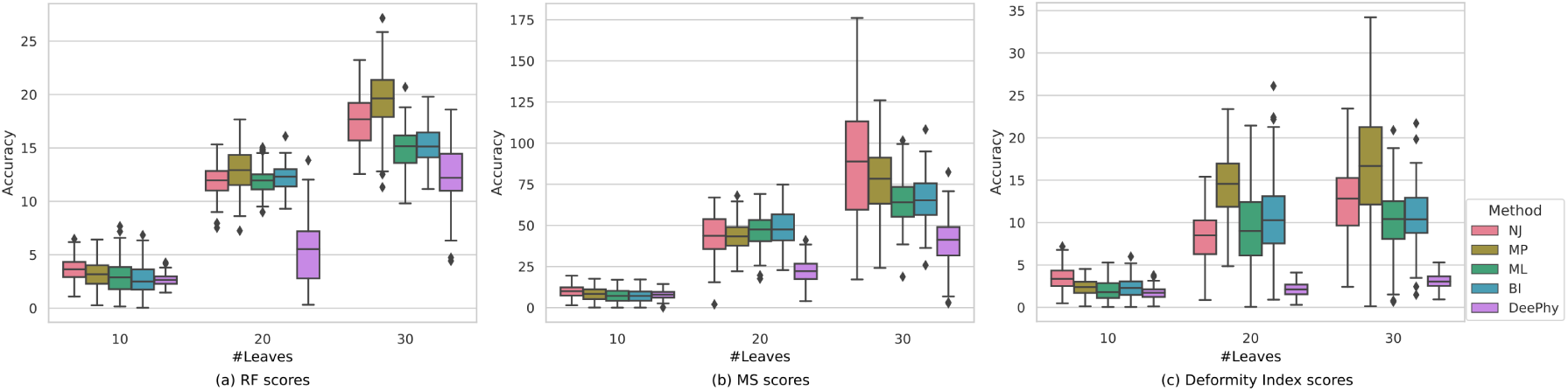
The (a) Robinson-Fould score, (b) Matching Split score, and (c) Deformity Index of the trees having various numbers of leaves. NJ = Neighbour Joining; MP = Maximum Parsimony; ML = Maximum Likelihood; BI = Bayesian Inference.

### 3.6 Performance on Biological dataset

From the previous results of DeePhy, it is observed that the performance of DeePhy is noticeably better than that of the traditional methods for data simulated under different conditions. Now, we demonstrate the evolutionary history by applying DeePhy on the biological dataset of Gadiformes. The list of species considered in this study is provided in the **Supplementary S3**. Since DeePhy is trained to derive the phylogenetic trees from the sequences of length ≤ 1356 bp, it is not suitable to reconstruct trees from the complete mitochondrial sequences at this stage. There exist few species whose several mitochondrial genes are also longer than 1, 356 bp (Férandon et al., 2010). Hence, we choose a biological dataset where the lengths of all the mitochondrial genes are within 1, 356 bp. Here, we use the trained model to derive the tree by considering each gene separately, which is referred to as the gene tree. The gene tree represents the evolution of the particular gene. The gene trees are provided in the **Supplementary S4**. Finally, we utilize the tool SuperTriplet to reconstruct the species tree from a set of gene trees. The species tree represents the evolution of the species. To understand the robustness of DeePhy, we apply our proposed bootstrapping technique and generate 100 bootstrap replicas from each gene separately. Since, we do not have the true reference tree topologies of the selected species of Gadiformes, we cannot assess the performance of the derived trees by using the RF and MS scores. Therefore, we compute DI of the derived trees which provides a measure of correctness based on the available hypotheses.

In this study, we consider 13 protein-coding genes of 20 species from the order Gadiformes. We derive the gene trees independently. Finally, by utilizing the SuperTriplet method, we amalgam them to reconstruct the phylogenetic tree. The Gadiformes dataset consists of eight major sub- families, such as Gadinae, Lotinae, Bregmacerotidae, Merlucciidae, Macrourinae, Trachyrincinae, Macrouroinae, and Bathygadinae. Though there are many hypotheses available regarding the re- lationships among the subfamilies of Gadiformes, few widely accepted hypotheses among them are listed below.

- Gadinae subfamily is monophyletic (von der Heyden and Matthee, 2008; Teletchea et al., 2006; Roa-Vaŕon and Ortí, 2009; Shi et al., 2016).
- Subfamily Lotinae is the sister group of Gadinae. Hence, both of them together form a monophyletic clade (Nelson, 1984; Fahay and Markle, 1984).
- Bregmacerotidae and Merlucciidae both form the monophyletic clade with Gadinae and Loti- nae (Teletchea et al., 2006; Endo, 2002; Shi et al., 2016; von der Heyden and Matthee, 2008).
- Macrourinae is monophyletic (Gaither et al., 2016).
- Macrouroinae and Trachyrincinae together form monophyletic clade (Kriwet and Hecht, 2008).
- Macrourinae, Macrouroinae, Trachyrincinae, and Bathygadinae together are monophyletic (Kri- wet and Hecht, 2008).

In the tree derived by utilizing DeePhy, as shown in Fig. 9, we observe that the Gadinae subfamily is monophyletic and Lotinae is the sister of Gadinae. However, all the members of the Macrourinae subfamily do not support the monophyletic clade. To assess the performance of the derived tree, we compute DI score of the derived tree with respect to the well-established hypotheses. We observe that the DI score for the tree derived by utilizing DeePhy (DI=7.395) is significantly less than that of the other standard methods for phylogenetic tree reconstruction, such NJ (DI=23.519), MP (DI=16.372), ML (DI=24.500), and BI (DI=13.257). This phenomenon indicates that the trees derived from DeePhy corroborates the existing hypotheses significantly more than that of the conventional methods with an acceptable bootstrap supports (*>* 75%) at almost each clade. So here for each method, we derive the gene trees, which further amalgam to infer the final tree. So in this approach, inferring an accurate tree also signifies that the respective gene trees are also inferred accurately. Therefore, we can expect that DeePhy derives the gene trees accurately from the mitochondrial gene sequences. However, it requires separate studies to analyze each gene trees derived from DeePhy.

**Figure 9:**
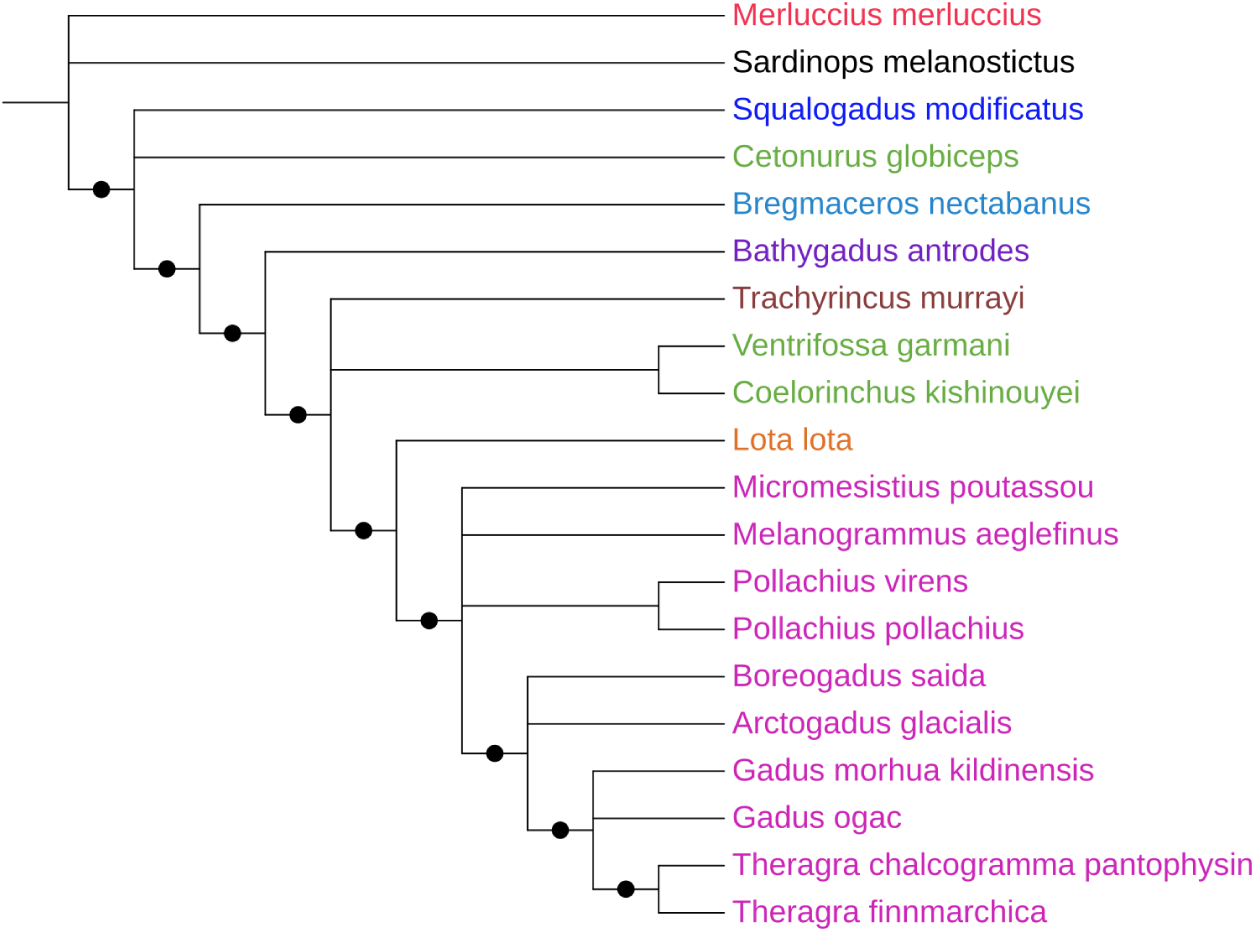
Phylogenetic tree of Gadiformes generated by utilizing DeePhy. There are different subfamilies, such as, Gadinae (pink), Lotinae (orange), Merlucciidae (red), Trachyrincinae (brown), Macrouroidae (blue), Bathygadinae (purple), Macrourinae (green), Bregmacerotidae (light blue). The outgroup is colored as black. The clade having the bootstrap score more than 75% are denoted as black dot.

## 4 Conclusion

In this study, we propose a convolutional neural network (CNN)-based model, called DeePhy, to infer the rooted triplet tree topology from a set of unaligned sequences. We utilize a technique Genomic Footprint (GFP) which is a biological semantics-based two dimensional representation scheme to represent an unaligned nucleotide sequence numerically. DeePhy takes the GFPs of three sequences as the input and provides a class level corresponding to a triplet tree topology as the output. After performing a numerous studies under different possible circumstances, the results exhibit that for a given set of three GFPs, DeePhy identifies the triplet tree topology quite well and it also has a number of desired qualities. Firstly, it performs more accurately than that of the conventional methods, like Neighbour Joining, Maximum Parsimony, Maximum Likelihood, and Bayesian Inference, in various scenarios. To identify the strength and the weakness of DeePhy more closely, we analyze the performances on the various narrowly defined branch-length subspaces. We observe that, for most of the branch-length zones DeePhy outperforms the conventional methods. For the rest of the zones, the performance of DeePhy is comparable with the best performing methods. When the performance of the traditional methods vary for different branch-length zones, the performance of DeePhy is found to be quite steady than the state-of-the-art methods among all the branch-length zones. To compute the robustness of the methods, here we also proposed a novel bootstrap technique to generate the replicas from the unaligned sequences. We are hopeful about the utility of the proposed bootstrap technique in the phylogenetic study especially when the alignment-free methods are considered for inferring trees. Based on the bootstrapping, we observe that it is decently robust and insensitive to the nucleotide sequence. We also extend this study by deriving the complete tree by utilizing a triplet tree amalgamation method. Both the Robinson-Fould score, the Matching Split score, and the Deformity Index demonstrate that the trees derived from our proposed method are significantly more accurate than that of the conventional methods. We train DeePhy on data of length 1, 356 bp. We observe that in various situations the trained model is able to determine the triplet tree topology accurately from the triplet sequences having length of ≤ 1356 bp each. We tried to apply DeePhy on the longer sequences by using a majority-consensus-based approach. The poor performance of this approach also indicates that the evolutionary signature may not be uniformly distributed within the genome sequences. However, we believe this limitation can be rectified by training DeePhy on the longer data. Since DeePhy shows encouraging performance when length of the sequence is ≤ 1356 bp, it has a profound utility to infer the gene trees more accurately than the state-of-the-art methods. Hence, we apply this method to reconstruct the gene trees from 13 mitochondrial genes of a biological dataset of gadiformes. We amalgam the gene trees to construct the final species tree of Gadiformes. Again, the Deformity Index of the derived phylogenetic tree of Gadiformes exhibits that the derived tree corroborates the existing hypotheses significantly more than that of the traditional methods. From the results, we expect that the gene trees derived from DeePhy are more accurate than that of the conventional methods. However, for a biological dataset, it is interesting to further analyze the evolution of each gene separately through the gene tree derived by utilizing DeePhy. Apart from that, the study can be extended to understand the gene flows among a set of species. Though DeePhy exhibits noticeable performance, but it only infers the tree topologies, not the associated branch lengths. So in future it will also be interesting to investigate the possibility of using deep learning-based methods to determine the branch-lengths in addition to the tree topology. Thus, although various practical improvements are still required, a deep learning-based approach may guide towards an exciting alternative way to address the problems related to tree inference and phylogenomic.

## Ethics declarations

### Conflict of interest

On behalf of all authors, the corresponding author states that there is no conflict of interest.

### Funding

This research did not receive any specific grant from funding agencies in the public, commercial or not-for-profit sectors.

### Informed Consent

All the data and personal information are collected from a public database. These data have been generated by various researchers since a decade.

### Authors’ contributions

AM and JM conceived the ideas; AM and JM designed methodology; AM collected the data; AM and JM analyzed the data; AM and JM led the writing of the manuscript. All authors contributed critically to the drafts and gave final approval for publication.

### Data Availability

A python-based deep learning framework, named ***DeePhy*** , for inferring triplet tree topology from three nucleotide sequences is freely available at http://www.facweb.iitkgp.ac.in/~jay/ DeePhy/DeePhy.html. The mirror of this program is also available at https://github.com/ aritramhp/DeePhy.

## Notes

### Competing Interest Statement

The authors have declared no competing interest.

