## Supplementary material for "DeePhy: A Deep Learning Model to Reconstruct Phylogenetic Tree from Unaligned Nucleotide Sequences": S1

### Grid Search

| # | Hidden layers | Fully connected layers | Pooling layer | Dropout | Learning rate | Accuracy |
|---|---|---|---|---|---|---|
| 1 | CNN(256) -> CNN(1024) -> CNN(256) | Fc(512) -> Fc(3) | MaxPool(1,3) | 0 | 0.01 | 67.17% (0.421) |
| 2 | CNN(256) -> CNN(1024) -> CNN(256) | Fc(512) -> Fc(3) | MaxPool(1,3) | 0 | 0.001 | 66.85% (0.324) |
| 3 | CNN(256) -> CNN(1024) -> CNN(512) | Fc(512) -> Fc(3) | MaxPool(1,3) | 0 | 0.001 | 66.06% (0.347) |
| 4 | CNN(256) -> CNN(1024) -> CNN(512) | Fc(512) -> Fc(3) | MaxPool(1,3) | 0 | 0.0001 | 67.41% (0.530) |
| 5 | CNN(256) -> CNN(1024) -> CNN(512) | Fc(512) -> Fc(3) | MaxPool(1,3) | 0 | 1E-05 | 66.98% (0.422) |
| 6 | CNN(64) -> CNN(128) -> CNN(64) | Fc(64) -> Fc(3) | MaxPool(1,3) | 0 | 0.001 | 68.90% (0.349) |
| 7 | CNN(64) -> CNN(512) -> CNN(64) | Fc(64) -> Fc(3) | MaxPool(1,3) | 0 | 0.001 | 68.56% (0.391) |
| 8 | CNN(64) -> CNN(1024) -> CNN(64) | Fc(64) -> Fc(3) | MaxPool(1,3) | 0 | 0.001 | 69.25% (0.455) |
| 9 | CNN(64) -> CNN(2048) -> CNN(64) | Fc(64) -> Fc(3) | MaxPool(1,3) | 0 | 0.001 | 69.35% (0.285) |
| 10 | CNN(64) -> CNN(128) -> CNN(128) -> CNN(64) | Fc(64) -> Fc(3) | MaxPool(1,3) | 0 | 0.001 | 71.45% (0.367) |
| 11 | CNN(64) -> CNN(128) -> CNN(128) -> CNN(128) -> CNN(64) | Fc(64) -> Fc(3) | MaxPool(1,3) | 0 | 0.001 | 69.42% (0.350) |
| 12 | CNN(64) -> CNN(128) -> CNN(256) -> CNN(128) -> CNN(64) | Fc(64) -> Fc(3) | MaxPool(1,3) | 0 | 0.001 | 69.96% (0.342) |
| 13 | CNN(64) -> CNN(128) -> CNN(512) -> CNN(128) -> CNN(64) | Fc(64) -> Fc(3) | MaxPool(1,3) | 0 | 0.001 | 68.96% (0.299) |
| 14 | CNN(64) -> CNN(128) -> CNN(512) -> CNN(64) | Fc(64) -> Fc(3) | MaxPool(1,3) | 0 | 0.001 | 70.96% (0.315) |
| 15 | CNN(64) -> CNN(128) -> CNN(128) -> CNN(64) -> CNN(64) | Fc(64) -> Fc(3) | MaxPool(1,3) | 0.25 | 1E-05 | 74.64% (0.336) |
| 16 | CNN(64) -> CNN(128) -> CNN(128) -> CNN(64) -> CNN(64) | Fc(64) -> Fc(3) | MaxPool(1,3) | 0.15 | 1E-05 | 78.2% (0.396) |
| 17 | CNN(64) -> CNN(128) -> CNN(128) -> CNN(64) -> CNN(64) | Fc(64) -> Fc(3) | MaxPool(1,3) | 0.1 | 1E-05 | 76.34% (0.359) |
| 18 | CNN(64) -> CNN(128) -> CNN(128) -> CNN(64) -> CNN(64) | Fc(64) -> Fc(3) | MaxPool(1,3) | 0.05 | 1E-05 | 75.72% (0.295) |
| 19 | CNN(64) -> CNN(128) -> CNN(128) -> CNN(128) -> CNN(64) | Fc(64) -> Fc(3) | MaxPool(1,3) | 0.1 | 1E-05 | 74.91% (0.258) |
| 20 | CNN(64) -> CNN(128) -> CNN(128) -> CNN(64) -> CNN(64) | Fc(64) -> Fc(3) | AvgPool(1,3) | 0.25 | 1E-05 | 75.51% (0.309) |
| 21 | CNN(64) -> CNN(128) -> CNN(128) -> CNN(64) -> CNN(64) | Fc(64) -> Fc(3) | AvgPool(1,3) | 0.15 | 1E-05 | 76.48% (0.358) |
| 22 | CNN(64) -> CNN(128) -> CNN(128) -> CNN(64) -> CNN(64) | Fc(64) -> Fc(3) | AvgPool(1,3) | 0.1 | 1E-05 | 79.65% (0.286) |
| 23 | CNN(64) -> CNN(128) -> CNN(128) -> CNN(128) -> CNN(64) | Fc(64) -> Fc(3) | AvgPool(1,3) | 0.15 | 1E-05 | 78.39% (0.382) |
| 24 | CNN(64) -> CNN(128) -> CNN(256) -> CNN(128) -> CNN(64) | Fc(64) -> Fc(3) | AvgPool(1,3) | 0.15 | 1E-05 | 77.92% (0.281) |
| 25 | CNN(64) -> CNN(128) -> CNN(256) -> CNN(128) -> CNN(64) | Fc(64) -> Fc(3) | AvgPool(1,3) | 0.15 | 1E-05 | 77.32% (0.219) |
| 26 | CNN(64) -> CNN(128) -> CNN(256) -> CNN(256) -> CNN(64) | Fc(64) -> Fc(3) | AvgPool(1,3) | 0.15 | 1E-05 | 73.96% (0.374) |
| 27 | CNN(64) -> CNN(256) -> CNN(256) -> CNN(256) -> CNN(64) | Fc(64) -> Fc(3) | AvgPool(1,3) | 0.15 | 1E-05 | 71.82% (0.391) |
