## Supplementary material for "DeePhy: A Deep Learning Model to Reconstruct Phylogenetic Tree from Unaligned Nucleotide Sequences": S2

### 1 MEGA Analysis Options (.mao) file for executing Neighbour Joining

```
[ MEGAinfo ]
ver                              = 10200331-x86_64 Linux
[ DataSettings ]
datatype                         = snNucleotide
containsCodingNuc                = False
MissingBaseSymbol                = ?
IdenticalBaseSymbol              = .
GapSymbol                        = -
Labelled Sites                   = All Sites
Labels to Include                =
[ ProcessTypes ]
ppInfer                          = true
ppNJ                             = true
[ AnalysisSettings ]
Analysis                         = Phylogeny Reconstruction
Scope                            = All Selected Taxa
Statistical Method               = Neighbor-joining
Phylogeny Test                   = ====================
Test of Phylogeny                = None
No. of Bootstrap Replications    = Not Applicable
Substitution Model               = ====================
Substitutions Type               = Nucleotide
Model/Method                     = Maximum Composite Likelihood
Substitutions to Include         = d: Transitions + Transversions
Rates and Patterns               = ====================
Rates among Sites                = Uniform Rates
Gamma Parameter                  = Not Applicable
Pattern among Lineages           = Same (Homogeneous)
Data Subset to Use               = ====================
Gaps/Missing Data Treatment      = Pairwise deletion
Site Coverage Cutoff (%)         = Not Applicable
System Resource Usage            = ====================
Number of Threads                = 1
Has Time Limit                   = False
Maximum Execution Time           = -1
```

#### 2 MEGA Analysis Options (.mao) file for executing Maximum Parsimony

```
[ MEGAinfo ]
ver                                  = 10180807-x86_64 Linux
[ DataSettings ]
datatype                             = snNucleotide
containsCodingNuc                    = False
MissingBaseSymbol                    = ?
IdenticalBaseSymbol                  = .
GapSymbol                            = -
[ ProcessTypes ]
ppInfer                              = true
ppMP                                 = true
[ AnalysisSettings ]
Analysis                             = Phylogeny Reconstruction
Statistical Method                   = Maximum Parsimony
Phylogeny Test                       = ====================
Test of Phylogeny                    = None
No. of Bootstrap Replications        = Not Applicable
Substitution Model                   = ====================
Substitutions Type                   = Nucleotide
Data Subset to Use                   = ====================
Gaps/Missing Data Treatment          = Use all sites
Site Coverage Cutoff (%)             = Not Applicable
Tree Inference Options               = ====================
MP Search Method                     = Subtree-Pruning-Regrafting (SPR)
No. of Initial Trees (random addition) = 10
MP Search level                      = 1
Max No. of Trees to Retain           = 100
Calculate Branch Lengths             = No
System Resource Usage                = ====================
Number of Threads                    = 1
Has Time Limit                       = False
Maximum Execution Time               = -1
```

### 3 MEGA Analysis Options (.mao) file for executing Maximum Likelihood

```
[ MEGAinfo ]
ver                               = 10180807-x86_64 Linux
[ DataSettings ]
datatype                          = snNucleotide
containsCodingNuc                 = False
MissingBaseSymbol                 = ?
IdenticalBaseSymbol               = .
GapSymbol                         = -
[ ProcessTypes ]
ppInfer                           = true
ppML                              = true
[ AnalysisSettings ]
Analysis                          = Phylogeny Reconstruction
Statistical Method                = Maximum Likelihood
Phylogeny Test                    = ====================
Test of Phylogeny                 = None
No. of Bootstrap Replications     = Not Applicable
Substitution Model                = ====================
Substitutions Type                = Nucleotide
Model/Method                      = General Time Reversible model
Rates and Patterns                = ====================
Rates among Sites                 = Gamma Distributed (G)
No of Discrete Gamma Categories   = 5
Data Subset to Use                = ====================
Gaps/Missing Data Treatment       = Use all sites
Site Coverage Cutoff (%)          = Not Applicable
Tree Inference Options            = ====================
ML Heuristic Method               = Nearest-Neighbor-Interchange (NNI)
Initial Tree for ML               = Make initial tree automatically (Default
    - NJ/BioNJ)
Branch Swap Filter                = None
System Resource Usage             = ====================
Number of Threads                 = 3
Has Time Limit                    = False
Maximum Execution Time            = -1
```

### 4 Parameter file for executing MrBayes

```
set autoclose=yes nowarn=yes
lset nst=6 rates=invgamma
prset statefreqpr = fixed(equal)
mcmc nruns=1 ngen=20000 samplefreq=1000
sump
sumt
quit
```
