## Supplementary material for "DeePhy: A Deep Learning Model to Reconstruct Phylogenetic Tree from Unaligned Nucleotide Sequences": S3

**Lengths of genes of each species**

| | Acc no. | ND1 | ND2 | ND3 | ND4 | ND4L | ND5 | ND6 | COI | COII | COIII | ATP8 | ATP6 | cytB | mtDNA |
|---|---|---|---|---|---|---|---|---|---|---|---|---|---|---|---|
| *Arctogadus glacialis* | AM919429 | 975 | 1045 | 349 | 1381 | 297 | 1839 | 522 | 1551 | 691 | 786 | 168 | 683 | 1141 | 16644 |
| *Bathygadus antrodes* | AP008988 | 975 | 1042 | 349 | 1381 | 297 | 1839 | 519 | 1551 | 691 | 785 | 162 | 683 | 1141 | 17596 |
| *Boreogadus saida* | AM919428 | 975 | 1045 | 349 | 1381 | 297 | 1839 | 522 | 1551 | 691 | 786 | 168 | 683 | 1141 | 16745 |
| *Bregmaceros nectabanus* | AP004411 | 969 | 1047 | 349 | 1374 | 297 | 1842 | 522 | 1557 | 691 | 784 | 168 | 681 | 1138 | 16030 |
| *Cetonurus globiceps* | KF751382 | 972 | 1050 | 349 | 1381 | 297 | 1839 | 522 | 1543 | 685 | 785 | 165 | 684 | 1140 | 17137 |
| *Coelorinchus kishinouyei* | AP002929 | 972 | 1047 | 349 | 1381 | 297 | 1845 | 522 | 1546 | 685 | 785 | 168 | 683 | 1140 | 15942 |
| *Gadus morhua kildinensis* | AM489716 | 975 | 1047 | 351 | 1386 | 297 | 1839 | 522 | 1551 | 699 | 786 | 168 | 684 | 1140 | 16654 |
| *Gadus ogac* | NC012323 | 975 | 1047 | 351 | 1386 | 297 | 1839 | 522 | 1551 | 699 | 786 | 168 | 684 | 1161 | 15564 |
| *Lota lota* | AP004412 | 975 | 1045 | 349 | 1381 | 297 | 1839 | 522 | 1551 | 691 | 785 | 168 | 683 | 1141 | 16527 |
| *Melanogrammus aeglefinus* | NC_007396 | 975 | 1045 | 349 | 1378 | 297 | 1839 | 522 | 1551 | 691 | 786 | 168 | 684 | 1141 | 16585 |
| *Merluccius merluccius* | NC_015120 | 975 | 1045 | 349 | 1381 | 297 | 1839 | 564 | 1551 | 691 | 786 | 168 | 684 | 1141 | 17087 |
| *Micromesistius poutassou* | FR751401 | 975 | 1045 | 349 | 1381 | 297 | 1839 | 522 | 1551 | 691 | 786 | 168 | 682 | 1141 | 16573 |
| *Pollachius pollachius* | NC_015097 | 975 | 1045 | 349 | 1381 | 297 | 1839 | 522 | 1551 | 691 | 786 | 168 | 682 | 1141 | 16539 |
| *Pollachius virens* | FR751399 | 975 | 1045 | 349 | 1381 | 297 | 1839 | 522 | 1551 | 691 | 786 | 168 | 682 | 1141 | 16556 |
| *Sardinops melanostictus* | NC_002616 | 975 | 1046 | 349 | 1381 | 297 | 1836 | 519 | 1551 | 691 | 785 | 168 | 683 | 1141 | 16881 |
| *Squalogadus modificatus* | AP008989 | 975 | 1045 | 351 | 1381 | 297 | 1842 | 522 | 1557 | 691 | 785 | 168 | 683 | 1141 | 16550 |
| *Theragra chalcogramma* | AB182305 | 975 | 1047 | 351 | 1386 | 297 | 1839 | 522 | 1551 | 699 | 786 | 168 | 684 | 1161 | 16571 |
| *Theragra finnmarchica* | AM489718 | 975 | 1047 | 351 | 1386 | 297 | 1839 | 520 | 1551 | 691 | 786 | 168 | 684 | 1141 | 16571 |
| *Trachyrincus murrayi* | AP008990 | 975 | 1045 | 349 | 1381 | 297 | 1839 | 522 | 1551 | 691 | 785 | 168 | 683 | 1141 | 16677 |
| *Ventrifossa garmani* | AP008991 | 972 | 1050 | 349 | 1381 | 297 | 1839 | 522 | 1548 | 685 | 785 | 165 | 683 | 1140 | 17230 |
