## Supplementary material for "DeePhy: A Deep Learning Model to Reconstruct Phylogenetic Tree from Unaligned Nucleotide Sequences": S4

### Gene trees derived by utilizing DeePhy

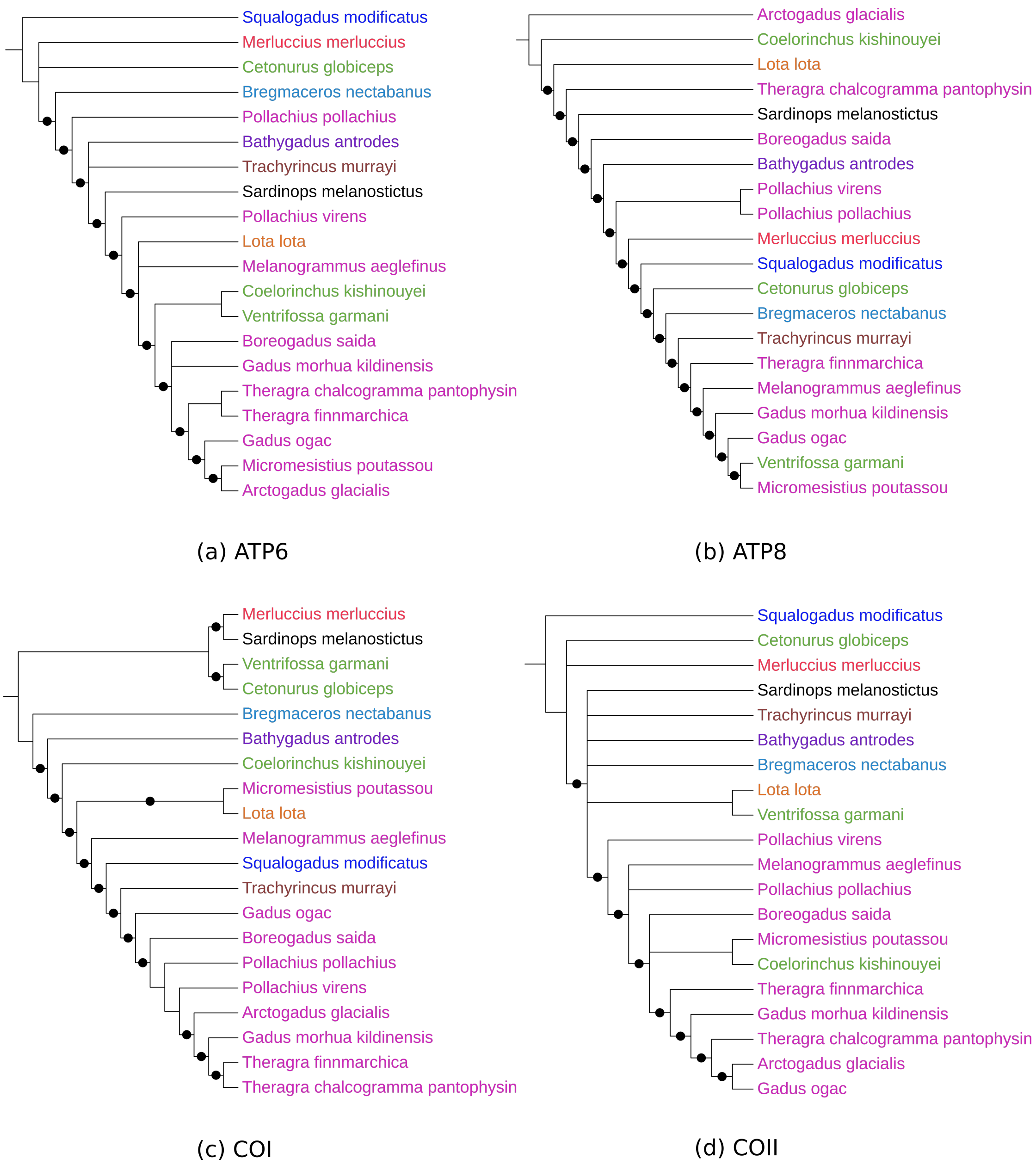


**Figure 1:** Gene trees derived by considering (a) ATP6, (b) ATP8, (c) COI, and (d) COII genes of Gadiformes generated by utilizing DeePhy. There are different subfamilies, such as, Gadinae (pink), Lotinae (orange), Merlucciidae (red), Trachyrincinae (brown), Macrouroidae (blue), Bathygadinae (purple), Macrourinae (green), Bregmacerotidae (light blue). The outgroup is colored as black. The clade having the bootstrap score more than 75% are denoted as black dot.

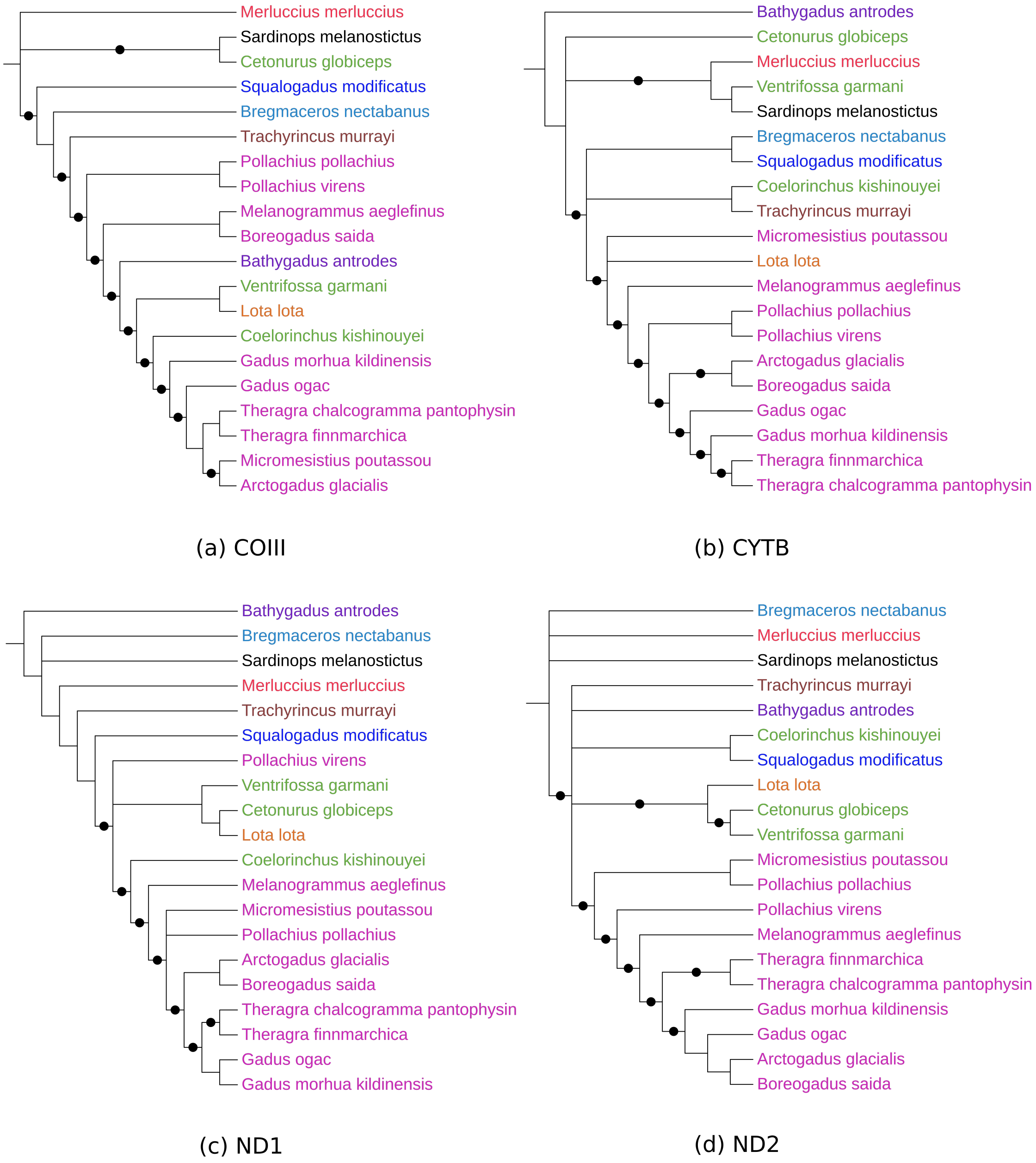


**Figure 2:** Gene trees derived by considering (a) COIII, (b) CYTB, (c) ND1, and (d) ND2 genes of Gadiformes generated by utilizing DeePhy. There are different subfamilies, such as, Gadinae (pink), Lotinae (orange), Merlucciidae (red), Trachyrincinae (brown), Macrouroidae (blue), Bathygadinae (purple), Macrourinae (green), Bregmacerotidae (light blue). The outgroup is colored as black. The clade having the bootstrap score more than 75% are denoted as black dot.

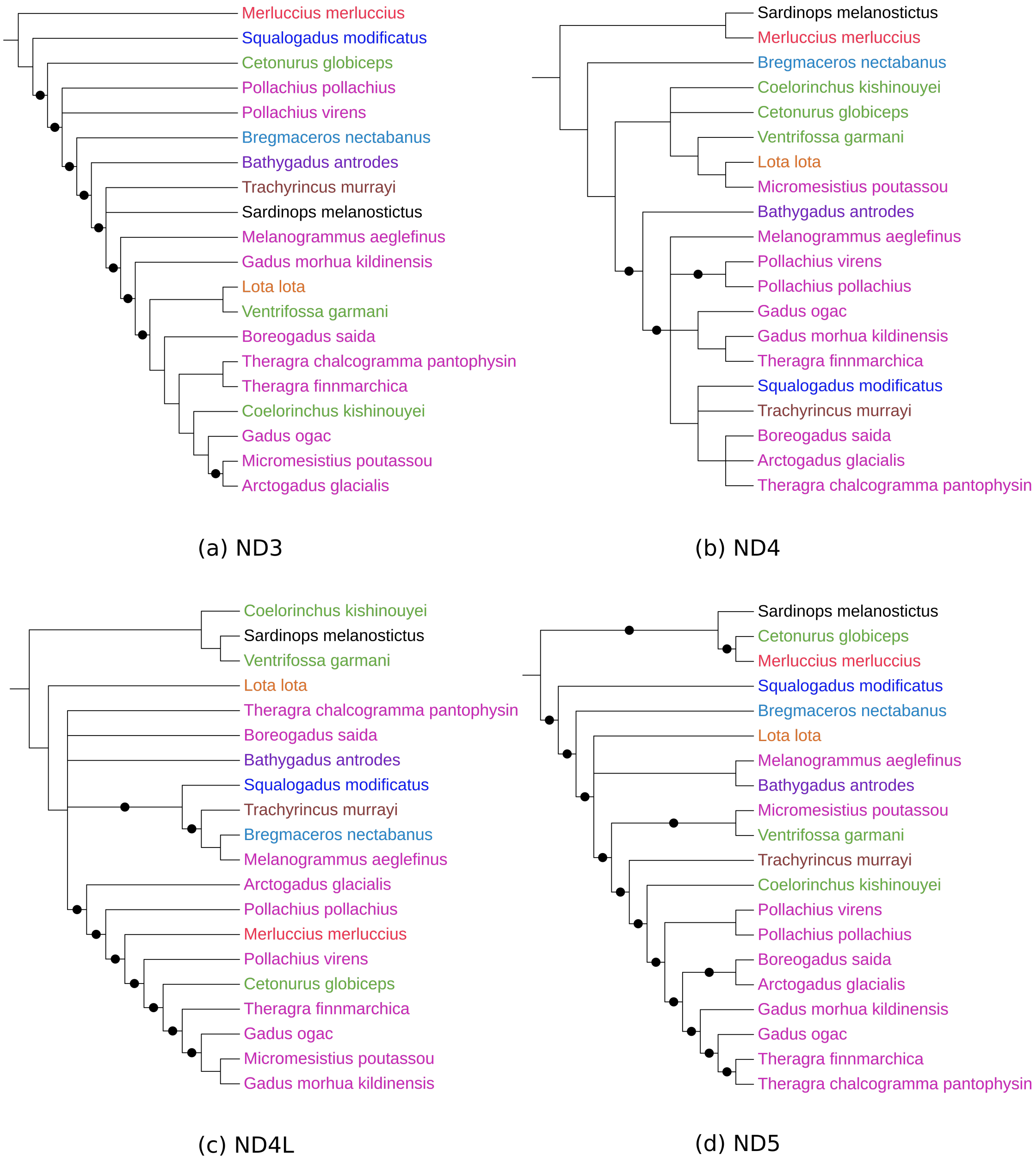


**Figure 3:** Gene trees derived by considering (a) ND3, (b) ND4, (c) ND4L, and (d) ND5 genes of Gadiformes generated by utilizing DeePhy. There are different subfamilies, such as, Gadinae (pink), Lotinae (orange), Merlucciidae (red), Trachyrincinae (brown), Macrouroidae (blue), Bathygadinae (purple), Macrourinae (green), Bregmacerotidae (light blue). The outgroup is colored as black. The clade having the bootstrap score more than 75% are denoted as black dot.

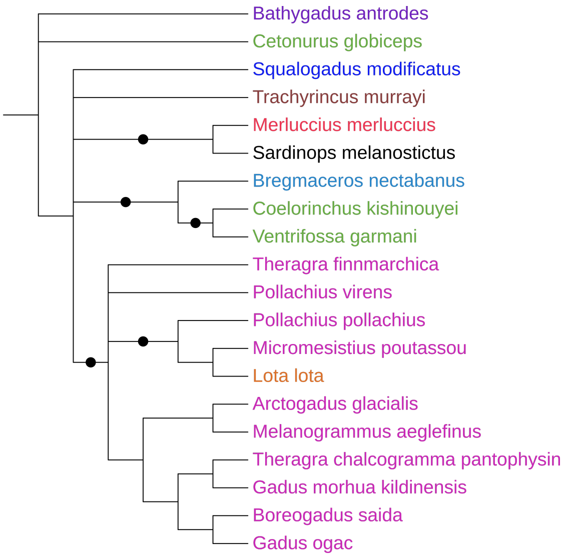


(a) ND6

**Figure 4:** Gene trees derived by considering (a) ND6 gene of Gadiformes generated by utilizing DeePhy. There are different subfamilies, such as, Gadinae (pink), Lotinae (orange), Merlucciidae (red), Trachyrincinae (brown), Macrouroidae (blue), Bathygadinae (purple), Macrourinae (green), Bregmacerotidae (light blue). The outgroup is colored as black. The clade having the bootstrap score more than 75% are denoted as black dot.
